# Arabidopsis Gluconolactonase, the First Enzyme Involved in Ascorbate Biosynthesis Localized in the Chloroplast Protects Plants from Light Stress

**DOI:** 10.1101/2024.02.22.578673

**Authors:** Jessica P. Yactayo-Chang, Nirman Nepal, Siddique I. Aboobucker, Karina Medina-Jiménez, Austin Wilkie, Thomas K. Teoh, Gwendolyn A. Wilson, Argelia Lorence

## Abstract

Vitamin C (L-ascorbic acid, AsA) is the most abundant water-soluble antioxidant in plants. Ascorbate scavenges free radicals, is an enzyme cofactor, and a donor and acceptor of electrons in the chloroplast. Ascorbate protects tissues against damage caused by reactive oxygen species (ROS) produced through normal metabolism or generated from stress. The inositol route to AsA involves four enzymes: *myo*-inositol oxygenase, glucuronate reductase, gluconolactonase (GNL), and L-gulono-1,4-lactone oxidase. The third enzyme, GNL, has been characterized in rat and bacteria but not in plants. Eighteen putative GNLs were identified in Arabidopsis, one of which, *AtGNL,* is interesting because it possesses a chloroplastic signal peptide. Plastids can accumulate up to 50 m M As A but until now no chloroplastic AsA biosynthetic genes have been described. This study includes the characterization of the first plant GNL enzyme *in vitro* and *in planta*. A knockout on this gene had lower foliar As A and stunted growth compared to controls. The functional gene restored the phenotype of the knockout, and those restored plants had higher AsA content, enhanced photosynthetic capacity, and higher seed yield. These results highlight the importance of *At*GNL in As A formation and in maintaining a healthy redox balance in the leaves particularly under low light stress.

## Introduction

Photosynthesis occurs in specialized organelles called chloroplasts that contain chlorophyll, which absorb light energy to produce ATP, fix atmospheric CO_2_ into sugars, and produce oxygen (Rustchow et al., 2008; Venkatasalam, 2012), Apart from photosynthesis, plastids are involved in other several specialized functions that include synthesis of amino acids, purine, pyrimidine bases, isoprenoids, tetrapyrroles, and the lipid components of their own membranes, followed by processing folding, and assembly by various chaperone systems (Peltier et al., 2006). Chloroplasts need considerable protein import from the cytosol and control nuclear gene expression indirectly by metabolites, ROS and other cellular processes (Pogson et al., 2008; Pfannschmidtm, 2010).

Reactive oxygen species are produced in different cell compartments. The major ROS production site is the apoplast due to the photosynthetic electron transport. Apoplastic ROS act as a signal and control primary and specialized metabolism in plants mediating response in polarized cell growth, stresses defense, cell-to-cell communication, and light dependent photosynthesis regulation. Ascorbate reduces ROS produced during the photo-respiration process (Foyer et al., 2019). Ascorbate plays important roles in systemic and local signaling pathways involved in growth and defense of plants (Córdoba-Pedregosa et al., 1996), and NO-mediated delaying senescence in *Oncidium* species (Kumar et al., 2016). Chloroplasts are a major source of ROS in green plant tissues. Light absorption creates oxidative stress due the formation of ROS including singlet oxygen (^1^O_2_), superoxide (O_2_^-^) and hydrogen peroxide (H_2_O_2_) (Oelze et al., 2008). Under high light, the electron flow through the photosynthetic chain overcomes the passage of electrons from ferredoxin to several reductases, and this causes an over-reduction of the plastoquinone and cytochrome b complex. Thus, during a day with high irradiance, plants are under constant oxidative stress (Oelze et al., 2008). Light/dark cycles are probably the most important signals that regulate plant development. Among the chief defense mechanisms that allow plants to cope with environmental stress situations is the ascorbate-glutathione cycle, a complex metabolic pathway in which a variety of photochemical and enzymatic steps are involved. Ascorbate is essential to detoxify H_2_O_2_ produced during the Mehler reaction, which is formed by dismutation of O_2_^-^ and can be regenerated via the AsA-glutathione cycle to counteract O_2_^-^ (Halliwell and Foyer, 1976; Foyer and Noctor, 2000; Munné-Bosch and Alegre, 2002; Talla et al., 2011). The demand for As A in these reactions increases at higher light intensities, when formation of ROS is enhanced. Experimental evidence suggests the existence of an effective signaling network between the chloroplasts and the mitochondria that involves ROS and antioxidants (Foyer and Noctor, 2003; Noctor et al., 2007). The different light/dark conditions are currently one of the major challenges in plant research to improve crop productivity under a changing global climate.

Ascorbate is present in all plants although its concentration varies greatly. Ascorbate can protect tissues against damage caused by ROS produced through normal oxygenic metabolism or generated from biotic, abiotic stress, and is strongly associated with photosynthesis and respiration. Other essential roles of As A is the modulation of processes such as lignification, cell division, cell elongation, the hypersensitive response, tolerance to stresses, and senescence in plants (Smirnoff and Wheeler, 2000; Barth et al., 2004; Pavet et al., 2005). In addition, As A controls flowering time through phytohormones (Barth et al., 2004). Ascorbate occurs inside as well as outside the chloroplasts (Constable, 1963; Hall and Rao, 1999; Habermann, 2013), where it has been shown to accumulate at concentrations up to 50 m M (Hall and Rao, 1999); this represents about 25 - 30% of the total As A in a cell (Horemans et al., 2000). All known As A biosynthetic enzymes reside in compartments other than the chloroplasts, and therefore it is currently unknown how this organelle is able to accumulate such high As A concentrations. Ascorbate was at one time considered to be a necessary component of the photosynthetic phosphorylation system (Arnon 1959) however is now considered important in providing a protective role in preventing inactivation of essential components of the chloroplasts (Pintó-Marijuan and Munné-Bosch, 2014).

The biosynthetic pathway for vitamin C in animals was elucidated in the early 1950s (Ishikawa et al., 2006). Evidence obtained during the last 25 years indicates that there are four pathways that lead to AsA synthesis in plants (Figure 1). These routes are the D-mannose/L-galactose (Wheeler et al., 1998), L-gulose (Wolucka and Van Montagu 2003), D-galacturonate (Agius et al., 2003), and *myo*-inositol (Lorence et al., 2004) pathways. The *myo*-inositol route involves four enzymes, starting from the oxidation of *myo*-inositol to D-glucuronic acid and reduction to L-gulonic acid and to L-gulono-1,4-lactone, and further oxidation to As A. These conversions are catalyzed by *myo*-inositol oxygenase (MIOX), glucuronate reductase (GlcUR), gluconolactonase (GNL), and L-gulono-1,4-lactone oxidase (GuILO), respectively. We have reported the biochemical characterization of MIOX and GuILO Arabidopsis isoforms (Lorence et al., 2004; Aboobucker et al., 2017). Using manual and digital phenotyping we have also shown that constitutive expression of *At*MIOX, and *At*GulLO and rat rGulLO leads to Arabidopsis plants with high AsA content, enhanced aerial and root tissues and tolerance to multiple abiotic stresses including heat, cold, salt and environmental pollutants (Lisko et al., 2013; Lisko et al., 2014; Yactayo-Chang et al., 2018).

**Figure 1.**
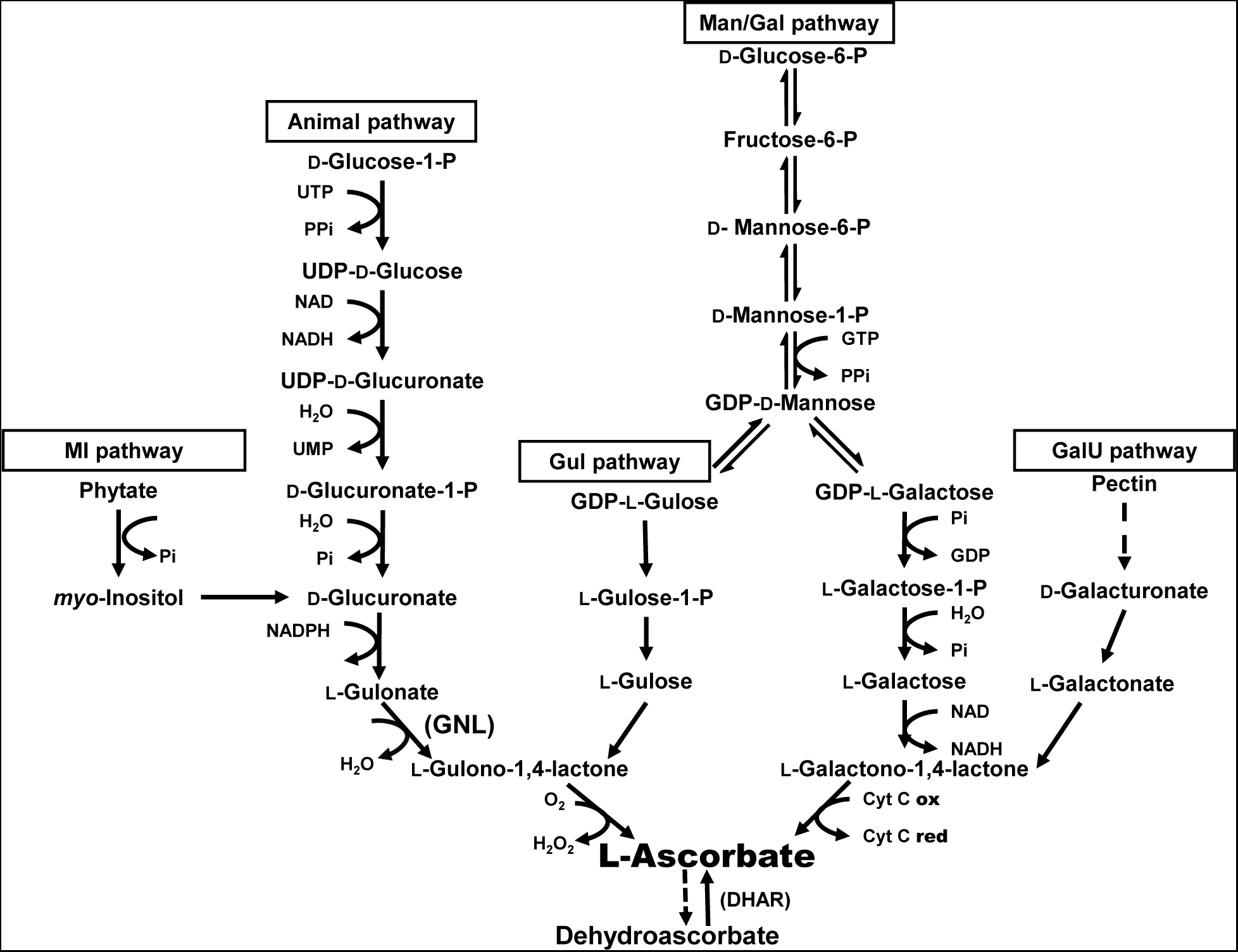
The ascorbate metabolic network. There is a single biosynthetic pathway for L-ascorbic acid (AsA) in animals, while four distinct routes lead its formation in plants: *myo*-inositol, L-gulose, D-mannose/L-galactose, and D-galacturonate pathways. The enzyme of interest in this work, gluconolactonase (GNL) that participates in the *myo*-inositol pathway is shown in bold letters.

Our next goal was to identify a functional gluconolactonase (GNL, EC 3.1.1.17) in Arabidopsis. Gluconolactonases have been characterized in *Rattus norvegicus* (Kondo et al., 2006), *Euglena gracilis* (Ishikawa et al., 2008), *Pseudomonas aeruginosa* (Tarighi et al., 2008), *Xanthomonas campestri* (Chen et al., 2008), *Gluconobacter oxydans* (Shinagawa et al., 2009) and *Homo sapiens* (Aizawa et al., 2013), but not in plants. Gene sequences of well characterized GNLs from rat and bacteria were aligned and compared to the *Arabidopsis* genome. This resulted in the identification of 18 putative *GNL* candidate genes (Table 1). T-DNA knockouts of putative GNLs were screening looking for low AsA lines to identify true GNLs enzymes in Arabidopsis. Bioinformatics analysis showed that one of the SALK lines with low AsA encodes a protein that possesses a chloroplastic signal peptide. In this work we demonstrated that At1g56500 encodes a functional GNL enzyme. Knocking down this gene leads to plants with low AsA, and the functional gene is able to restore the AsA content of the knockouts. Physiological assays show that AtGNL is able to protects plants from light stress. Over-expression of AtGNL enhances AsA content in plants and leads to over-expressers that grow faster, accumulate more biomass, posses enhanced photosynthetic efficiency, and higher seed yield.

**Table 1.** Putative glucuronolactonases (GNLs) in Arabidopsis.

| N° | Locus | Knockout | Predicted Localization | Notes |
| --- | --- | --- | --- | --- |
| 1 | *At1g08470* | SALK_055631 | NA | Strictosidine synthase family protein |
| 2 | *At1g56500* | SALK_011623<br>SALK_026172 | Chloroplast | Haloacid dehalogenase-like hydrolase family |
| 3 | *At1g74000* | SALK_032305 | Cell wall | Encodes a protein similar to strictosidine synthase |
| 4 | *At1g74020* | SALK_001853 | Endomembrane system | Responds to JA; strictosidine synthase family protein |
| 5 | *At2g01410* | SALK_147211 | Endomembrane system | Expressed protein; AsA |
| 6 | *At2g16780* | SALK_001214 | NA | WD-40 repeat protein (MSI2), contains 5 WD-40 repeats (PF0400) |
| 7 | *At2g24130* | SALK_025037 | Endomembrane system | Leucine-rich repeat transmembrane protein kinase, putative |
| 8 | *At2g41290* | SALK_015649 | Endomembrane system | Strictosidine synthase family protein |
| 9 | *At2g41300* | SALK_009989 | Endomembrane system | Strictosidine synthase family protein |
| 10 | *At3g51420* | SALK_026466 | Endomembrane system | Strictosidine synthase family protein |
| 11 | *At3g51430* | SALK_055182 | Endomembrane system | Strictosidine synthase |
| 12 | *At3g51440* | SALK_057855 | Endomembrane system | Strictosidine synthase family protein |
| 13 | *At3g51450* | SALK_124438 | Endomembrane system | Strictosidine synthase family protein |
| 14 | *At3g57010* | SALK_091299 | Endomembrane system | Strictosidine synthase family protein |
| 15 | *At3g57020* | NA | Endomembrane system | Strictosidine synthase family protein |
| 16 | *At3g57030* | SALK_120135 | Cell wall | Strictosidine synthase family protein |
| 17 | *At3g59530* | SALK_033871 | Endomembrane system | Strictosidine synthase family protein |
| 18 | *At5g22020* | SALK_042799 | Endomembrane system | Strictosidine synthase family protein |

## RESULTS

### Putative Glucuronolactonase (GNLs) in Arabidopsis

Glucuronalactonase (GNL), the third enzyme in the MI pathway (Figure 1) has been characterized in rat, bacteria’s and other organism but not in plants. The *GNL* gene sequences of well characterized GNLs from rat and bacteria were aligned and compared to the Arabidopsis genome. This resulted in the identification of 18 knockout *GNL* candidate genes (Table 1). The foliar ascorbate content was measured in all available knockout lines to identify low AsA mutants. Bioinformatic analysis was performed and identified that one of the knockouts with low As A encodes a protein that possesses a chloroplastic signal peptide. In addition, microarray data available at Genevestigator (Zimmerman et al., 2004) showed that there is a suppression of the expression of this gene when plants are exposed to dark. To understand the role of the *At1g56500* (*At*GNL) gene in plant physiology, two knockouts lines (SALK_026172 and SALK_011623) with a T-DNA inserted into the *At*GNL gene were obtained from the Arabidopsis Biological Resources Center (ABRC). The site of the insertion of the T-DNA in each of these knockouts is illustrated in Figure 2A. The *At*GNL cDNA was amplified and subcloned into pBIB-Kan under the control of the cauliflower mosaic virus 35S promoter and the tobacco etch virus (TEV) enhancer. A 6X-HIS tag was added at the 5’ end of the cDNA to facilitate protein detection (Figure 2B-C).To confirm that indeed this gene encodes a protein residing in the chloroplast, *N. benthamiana* plants were infiltrated with the *At*GNL construct using an optimized *Agrobacterium*-mediated transient transformation method (Medrano et al., 2009). Chloroplasts were isolated from leaves using a chloroplast isolation kit (CP-ISO, Sigma). A Western blot developed with and anti-HIS antibody, confirmed that *At*GNL is indeed in the chloroplasts as illustrated in Figure 3.

**Figure 2.**
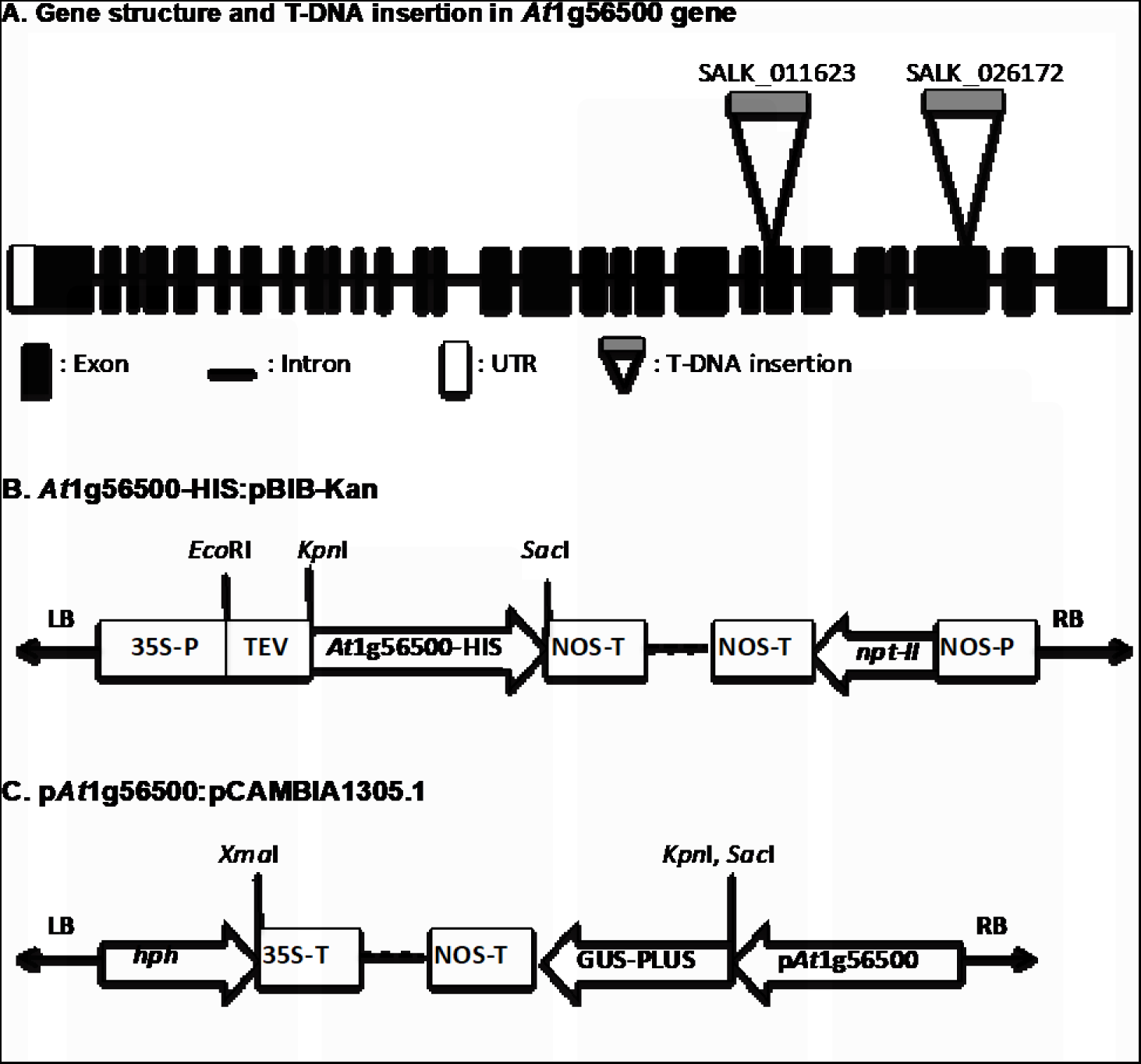
Constructs of interest. (A) Schematic representation of T-DNA insertion in SALK_011623 and SALK_026172 lines. (B) The *At*1g56500-6XHIS:pBIB-Kan construct containing *At*1g56500 (*At*GNL) with a six histidine (6X-HIS) tag and adjacent neomycin phosphotransferase II (*npt*II) selectable marker. (C) The p*At*1g56500:pCAMBIA1305.1 construct containing the *At*GNL promoter with the *GUS-PLUS* reporter gene and the hygromycin phosphotransferase (*hph*) selectable marker. NOS-P: promoter of nopaline synthase gene, 35S-T: terminator of the 35S cauliflower mosaic virus gene; TEV: tobacco etch virus translational enhancer; LB and RB: left and right T-DNA borders, respectively.

**Figure 3.**
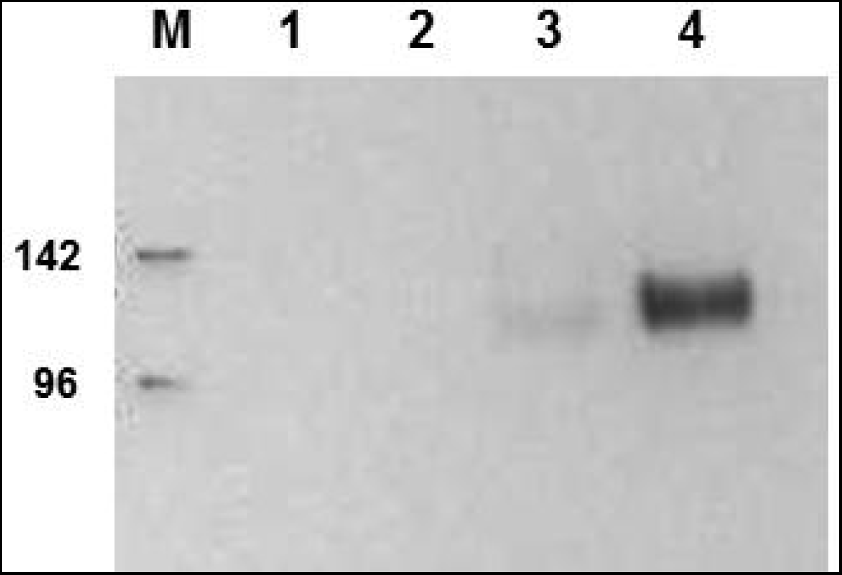
The recombinant *AtGNL* protein resides in the chloroplast. *At*GNL detection was done using Western blot and an anti-HIS antibody. M: Molecular weight marker, lane 1: empty vector fraction, lane 2: non-chloroplastic fraction, lane 3: chloroplastic fraction, lane 4: chloroplast fraction after purification by nickel affinity chromatography.

### Purification and Characterization of Recombinant *At*GNL

A *Nicotiana benthamiana*-based transient expression system was used to demonstrate the GNL activity *in vitro*. Plants were vacuum infiltrated with *A. tumefaciens* LBA4404 strain carrying the *At*GNL construct, proteins were separated by SDS-PAGE and *At*GNL was detected by Western blot. The optimal time for protein accumulation was at 48 h post infiltration (data not shown). Various buffers were tested to identify one that will yield the highest protein recovery. Of the tested buffers, one containing sodium phosphate, sodium metabisulfite and sodium chloride gave the best results (data not shown). The optimal concentration for washing and elution buffer with imidazole concentrations were 40 and 250 m M, respectively. Western blot results showed the presence of *At*GNL in the crude extract and flow through indicating the protein had a partial binding to the cation column, and the silver-stained gel indicates that the protein preparation contained mostly the protein of interest with a few minor contaminants (Figure 4).

**Figure 4.**
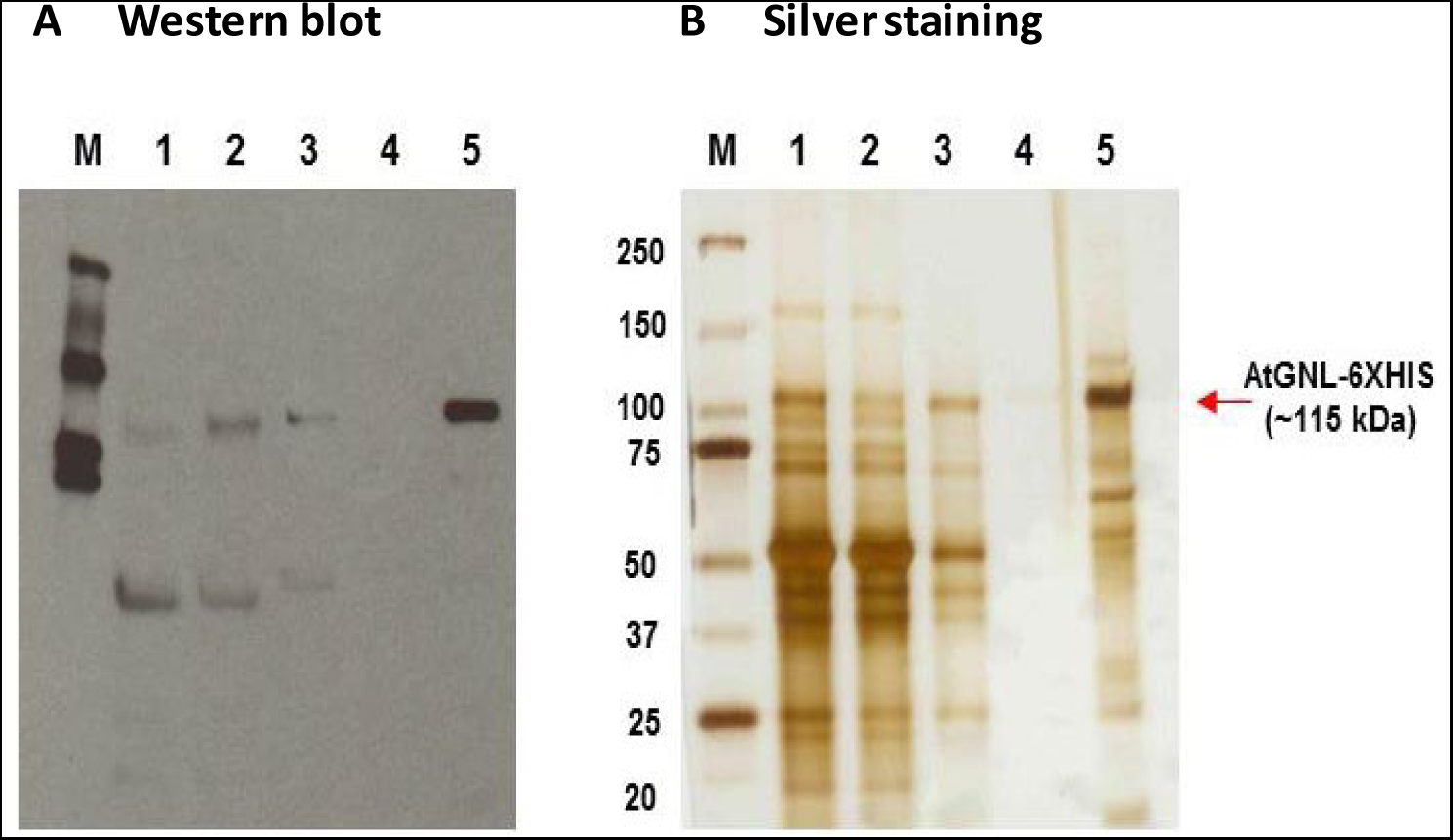
Purification of the *At*GNL-6XHIS;pBIB-Kan expressed in *Nicotiana benthamiana* leaves. (A) Western blot (B) Silver staining. M: marker, lane 1: crude extract, lane 2: flow through, lane 3: wash, lame 4: enzyme, lane 5: concentrated enzyme.

### Enzyme Activity of Gluconolactonase

Gluconolactonase catalyzes the hydrolysis of D-glucono-σ-lactone (D-GuIL) to D-gluconic acid (Ogawa et al., 2002). The lactonase activity was assayed *in vitro* based on the decrease in absorbance (405 nm) of the *p*-nitrophenol pH indicator that resulted from the enzymatic opening of the lactone ring when D-glucono-σ-lactone (D-GuIL) was used as substrate in presence of the *At*GNL as previously described (Hucho and Wallenfels, 1972). The enzymatic activity was assayed at 25°C with 10 m M PIPES pH 6.5, 5 m M D-GuIL, 75 µM MnCl_2_, 2.5 m M *p*-nitrophenol, and 30 µg of the purified enzyme (*At*GNL) in 1 mL of reaction (Ishikawa et al., 2008). Except for the D-GuIL, the recombinant *At*GNL did not exhibit activity with any of the substrates tested such as: L-galactono-ɤ-lactone, L-galactonic acid, L-gulono-ɤ-lactone and L-gulonic acid (Table 2).

**Table 2.** Comparison of *At*GNL with other well characterized GNLs.

| Species | pH | Cofactor | Specific activity<br>($\mu\text{mol min}^{-1} \text{mg}^{-1}$ of protein) | Km<br>(mM) | Vmax<br>($\mu\text{mol min}^{-1} \text{mg}^{-1}$ of protein) | Ref |
| --- | --- | --- | --- | --- | --- | --- |
| *Arabidopsis thaliana* | 6.0 | Mn<sup>2+</sup>, Mg<sup>2+</sup>, Zn<sup>2+</sup> | 10.54 | 3 | 38.7 | This work |
| *Rattus norvegicus* | 6.4 | Zn<sup>2+</sup> | 9 | 9.4 | 345 | Kondo *et al.*, 2006 |
| *Euglena gracilis* | 6.5 | Zn<sup>2+</sup> | 39 | 9.4 | 345 | Ishikawa *et al.*, 2008 |
| *Pseudomonas aeruginosa* | 7.2 | not mentioned | not mentioned | not mentioned | not mentioned | Tarighi *et al.*, 2008 |
| *Xanthomonas campestris* | 7.0 | Ca<sup>2+</sup> | not mentioned | 16.2 | 160 | Chen *et al.* 2009 |
| *Gluconobacter oxydans* | 6.0 | Ca<sup>2+</sup> | not mentioned | 2.5 | 25 | Shinagawa *et al.*, 2009 |
| *Homo sapiens* (SMP30)* | 6.4 | Ca<sup>2+</sup> | 25 | not mentioned | not mentioned | Aizawa *et al.*, 2013 |
| *Mus musculus* | 6.4 | Ca<sup>2+</sup> | 8.8 | not mentioned | not mentioned | Aizawa *et al.*, 2013 |

\*The human GNL a.k.a. senescence marker protein 30 or SMP30 is a protein whose tissue levels in the liver, kidney and lung decrease with aging.

The effect of the temperature on the *At*GNL activity was also determined (Figure 5A). The activity was highest at temperatures between 25°C and 35°C. Enzyme activity decreased when the temperature was increases to 40°C. In this study the *A. thaliana* GNL enzyme had a higher activity at pH 6.0, and the activity decreased by 4-fold when pH was increased to 6.3 (Figure 5B). To assess if the *At*GNL activity preferred a particular divalent ion, various cofactors were tested. In these experiments, no significant difference in GNL activity among the tested cofactors was observed (MnCl_2_, MgCl_2_, and ZnCl_2,_ Supplemental Figure 1). Increasing the substrate concentration elevated the rate of the reaction or enzyme activity. After testing different concentrations of substrate, we concluded that 3 m M of D-glucono-δ-lactone, was the most effective substrate concentration (Supplemental Figure 1).

**Figure 5.**
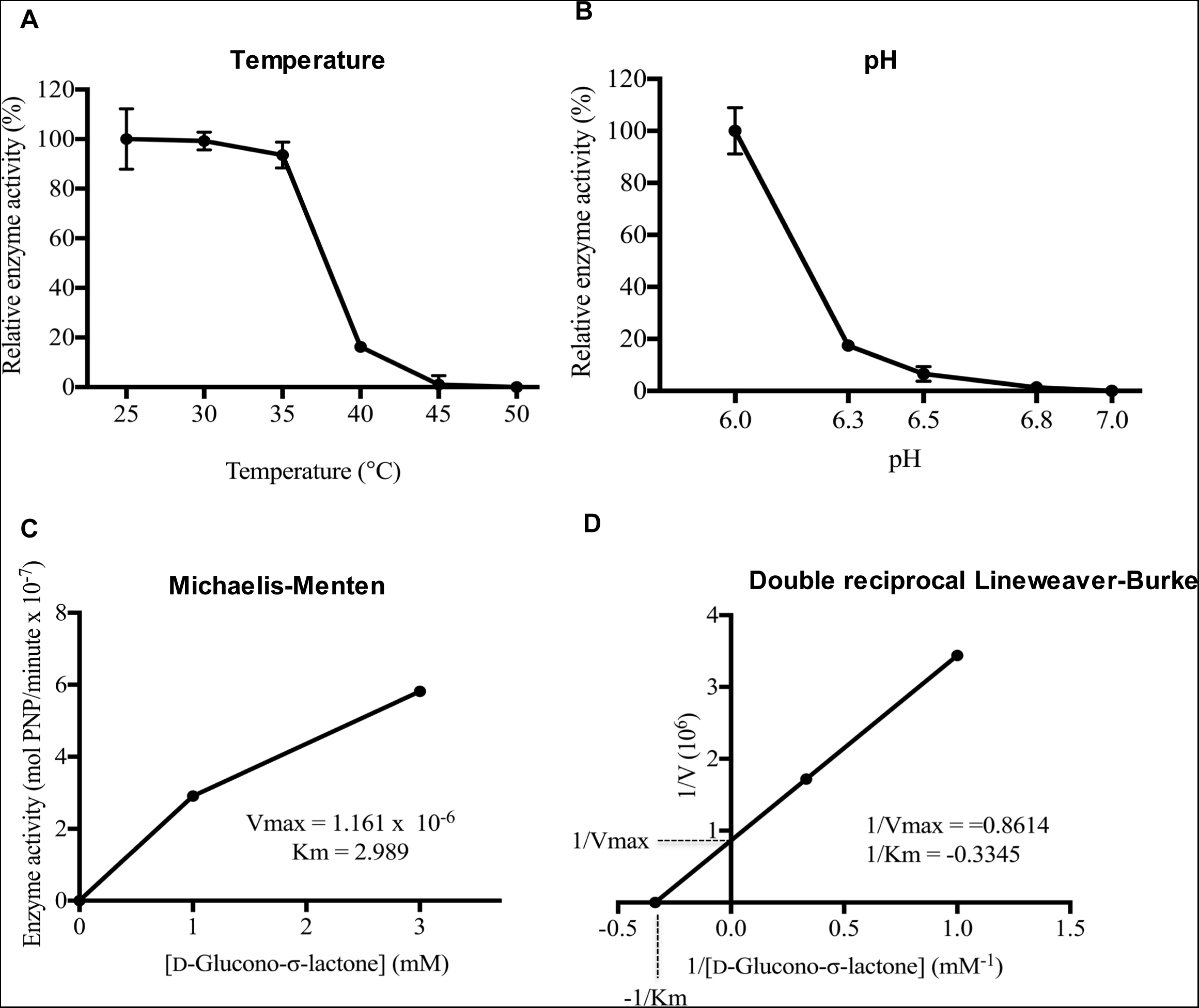
Effects of temperature and pH on *At*GNL enzyme activity. (A) pH effect on *At*GNL activity. (B) Temperature effect on *At*GNL activity. (C) Michaelis-Menten plot (D) Double reciprocal Lineweaver-Burke plot. Measurements were made in duplicate. Values are means ± SD.

Enzyme kinetic analysis was performed with D-GuIL at a concentration of 1 to 50 mM of substrate. The enzyme activity with 5 m M of D-GuIL at pH 6.0 was 10.54 µmol/min/mg of protein, V_max_ = 1.161 x 10^-6^ (38.7 µmol/min/mg of protein) and Km = 2.989 (Figure 5C-D). Table 3 summarizes the comparison between the kinetic parameters of the *At*GNL with the one of known GNLs. Based on these results the *G. oxidans* GNL is the most similar to the Arabidopsis GNL.

### Characterization of the Phenotype of Arabidopsis Lines

Primary transformants that were PCR positive (135) were screened to identify high AsA expressers. After four rounds of screening, 3 lines per group were selected for further analysis: over-expressers (WT+*At*GNL), restored 1 (SALK_026172+*At*GNL), and restored 2 (SALK_011623 + *At*GNL). Homozygous lines (T5), plants with 100% germination in the presence of antibiotic selection were developed for over-expresser (L60, L61, and L62), restored-1 lines (L89, L90, and L91), and restored-2 (L128, L129, and L130). Total foliar AsA content was measured in all homozygous lines. As illustrated in Figure 6 AtGNL constitutive expression leads to plants with elevated foliar As A in both the WT and T-DNA insertion line backgrounds. The phenotype of these homozygous lines was analyzed using a high throughout phenotyping instrument. Images were captured every two days from 16 to 26 days after germination. Representative images of homozygous *At*GNL lines and their controls are shown in Figure 7. As illustrated there is a strong correlation between higher foliar As A content, biomass, and project leaf area. Restored-1 (R3) and restored-2 (R5) lines had more biomass compared to their controls SALK_026172, SALK_011623, respectively, and those restored lines presented higher projected leaf area compared with controls.

**Figure 6.**
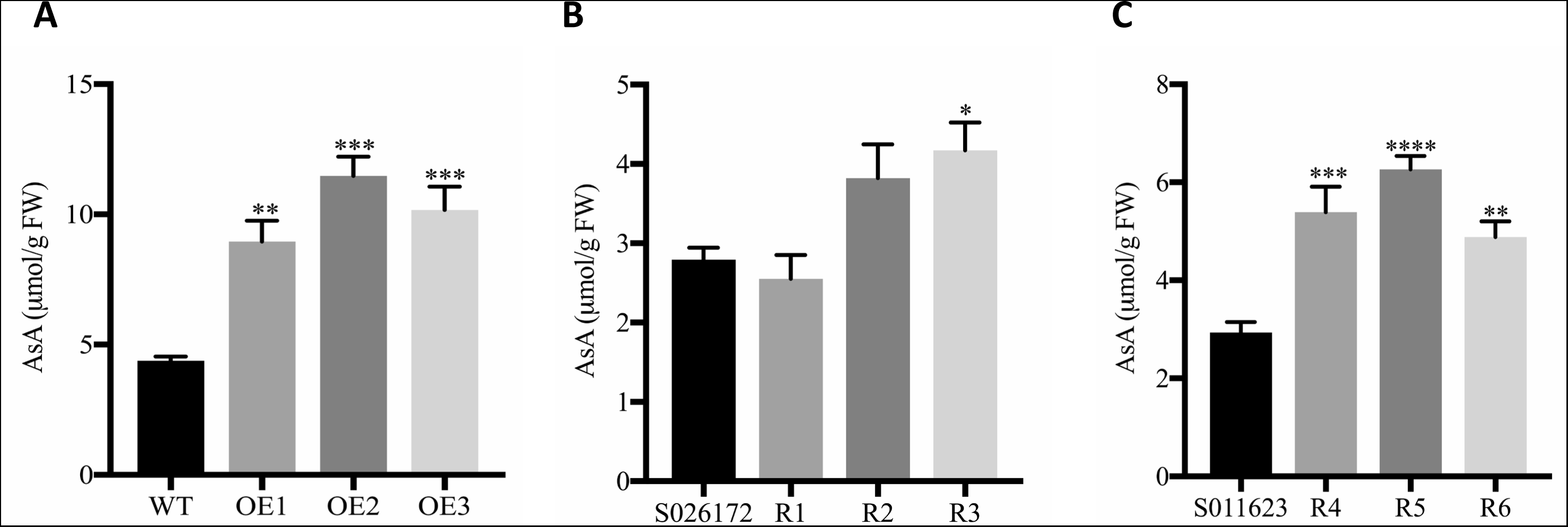
Total foliar ascorbate content of Arabidopsis lines of interest grown under normal light conditions. (A) *Arabidopsis thaliana* var. Columbia (Col-0, WT) and three *At*GNL over-expressers (OE1, OE2, OE3). (B) SALK_026172 T-DNA insertion line and three restored lines (R1, R2 and R3). (C) SALK_011623 T-DNA insertion line and three restored lines (R4, R5, R6). Asterisks indicate significant differences between controls and high As A lines as determined by Tukey’s posthoc multiple comparisons test, ****=p˂0.0001, ***=p˂0.0005, **=p˂0.0017, *=p˂0.0155. Values are means ± SEM (n=15).

**Figure 7.**
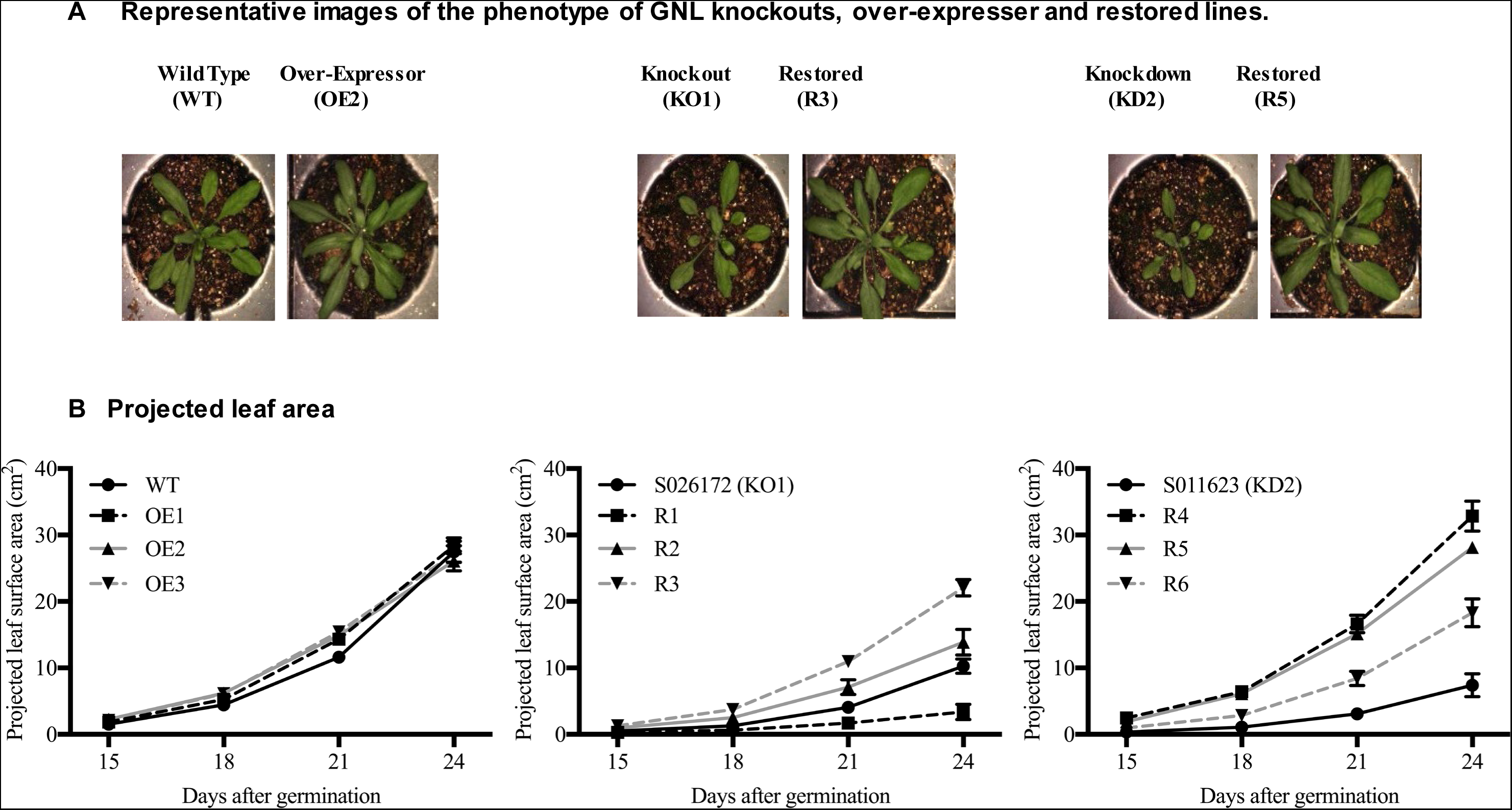
Phenotype of *At*GNL lines grown under normal conditions. (A) Representative images of *At*GNL lines acquire with the RGB camera. (B) Growth curves of *At*GNL lines compared with their respective controls. Values are means ± SEM (n=15).

RT-qPCR analysis was used to determine if SALK_026172 and SALK_011623 were true knockouts, and to confirm the increase AtGNL expression in the over-expressers and restored lines. As shown in Figure 8 the *GNL* transcript is significantly lower than the one in the WT only in SALK_026172, while is increased in the over-expressers and restored lines. From this point forward, we refer to SALK_026172 as knockout 1 (KO1) and to SALK_011623 as knockdown 2 (KD2).

**Figure 8.**
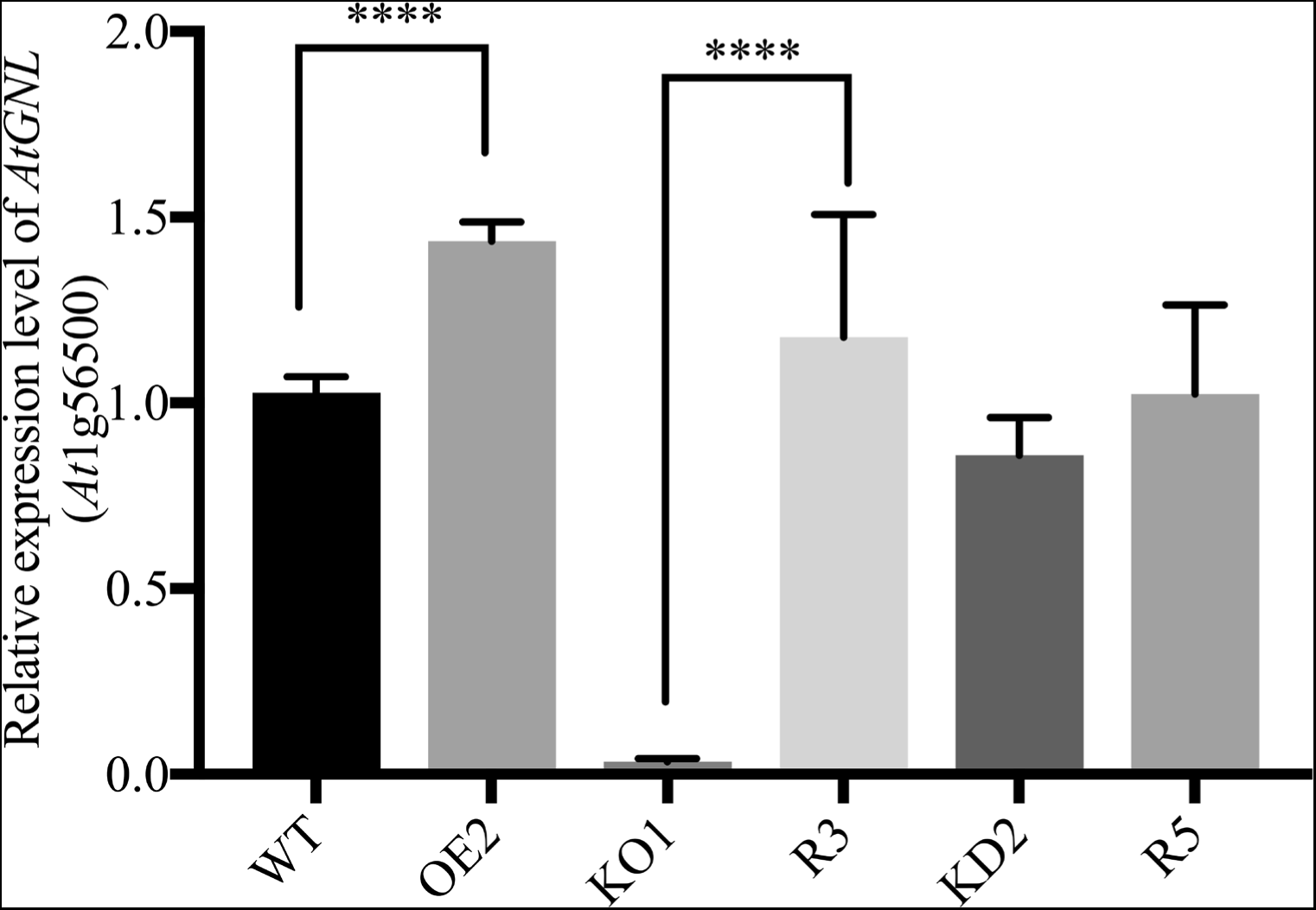
Expression of *At*GNL quantified by RT-qPCR. Student’s t-test was used to compare *At*1g56500 expression in *At*GNL lines with the respective controls. **** indicates significant differences at p˂ 0.0001 at 0.05 significance level. WT: wild type, OE: over-expresser, KO: knockout, KD: Knockdown, R: restored.

Based on these results, further studies were done only with the lines that had highest foliar AsA content, faster growth, higher biomass and projected leaf area. Over-expresser L61 (OE2) and the empty vector control (EV); restored-1 L100 (R3) and its control SALK_026172 (KO1), restored-2 L129 (R5), and its control SALK_011623 (KD2), and wild type (WT) control were selected to study the effect of low (35-110 µmol/m^2^/s), normal (110-350 µmol/m^2^/s), and high light (350-700 µmol/m^2^/s) conditions in a greenhouse. The projected leaf area results showed the same trend, where over-expressers and restored lines were bigger than their controls, with KO1 being the worst performer at all light conditions tested (Figure 9). The results indicate that the over-expresser and the restored lines had higher foliar As A than their respective controls growing under similar conditions (Figure 10). There is a strong correlation between foliar As A levels and higher projected leaf area in the high AsA lines compared with their respective controls.

**Figure 9.**
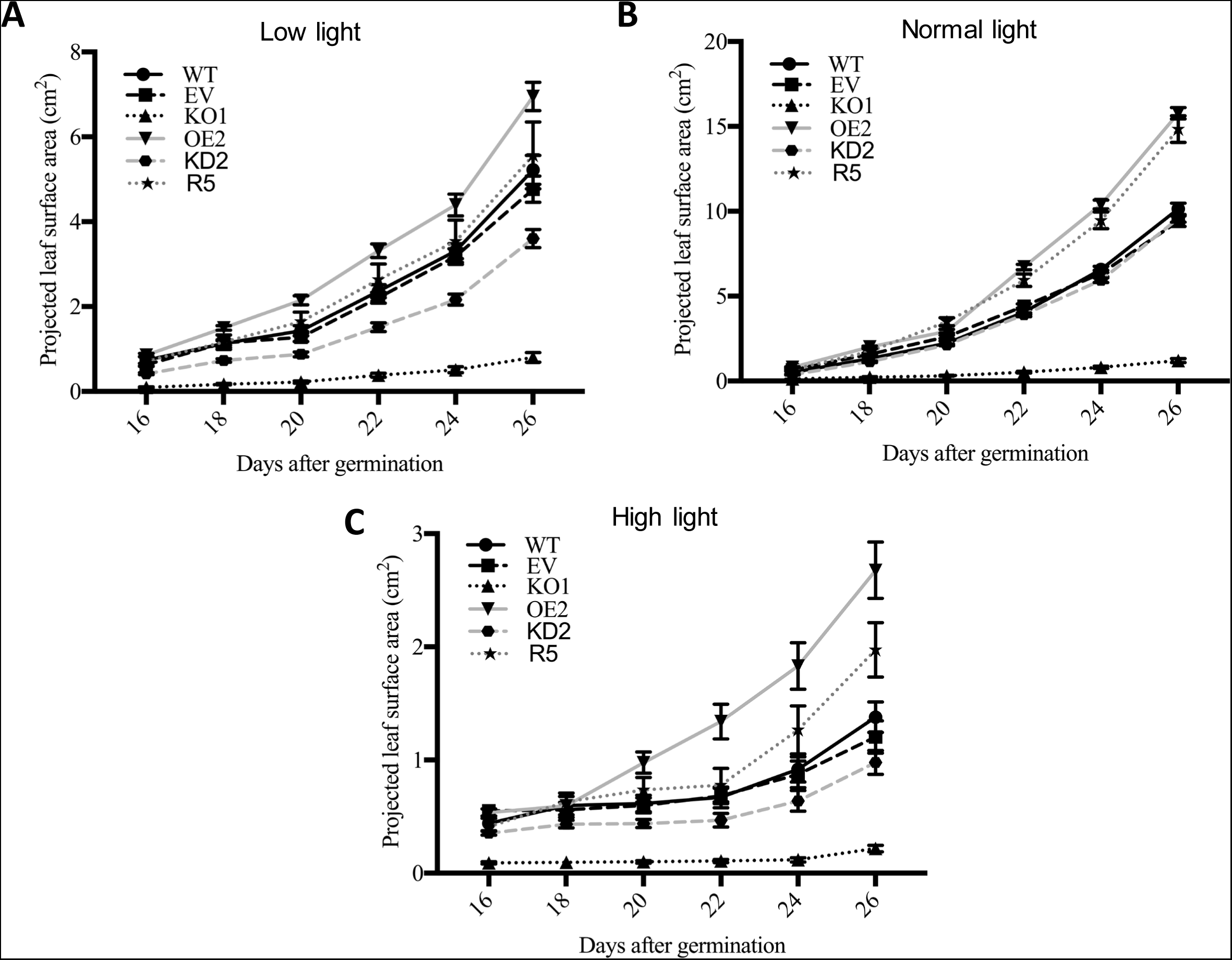
Growth curve of *At*GNL lines growing under (A) low light, (B) normal light, and (C) high light conditions. WT: wild type, EV: empty vector, OE: over-expresser, KO: knockout, R: restored. Values are means ± SEM (n=15).

**Figure 10.**
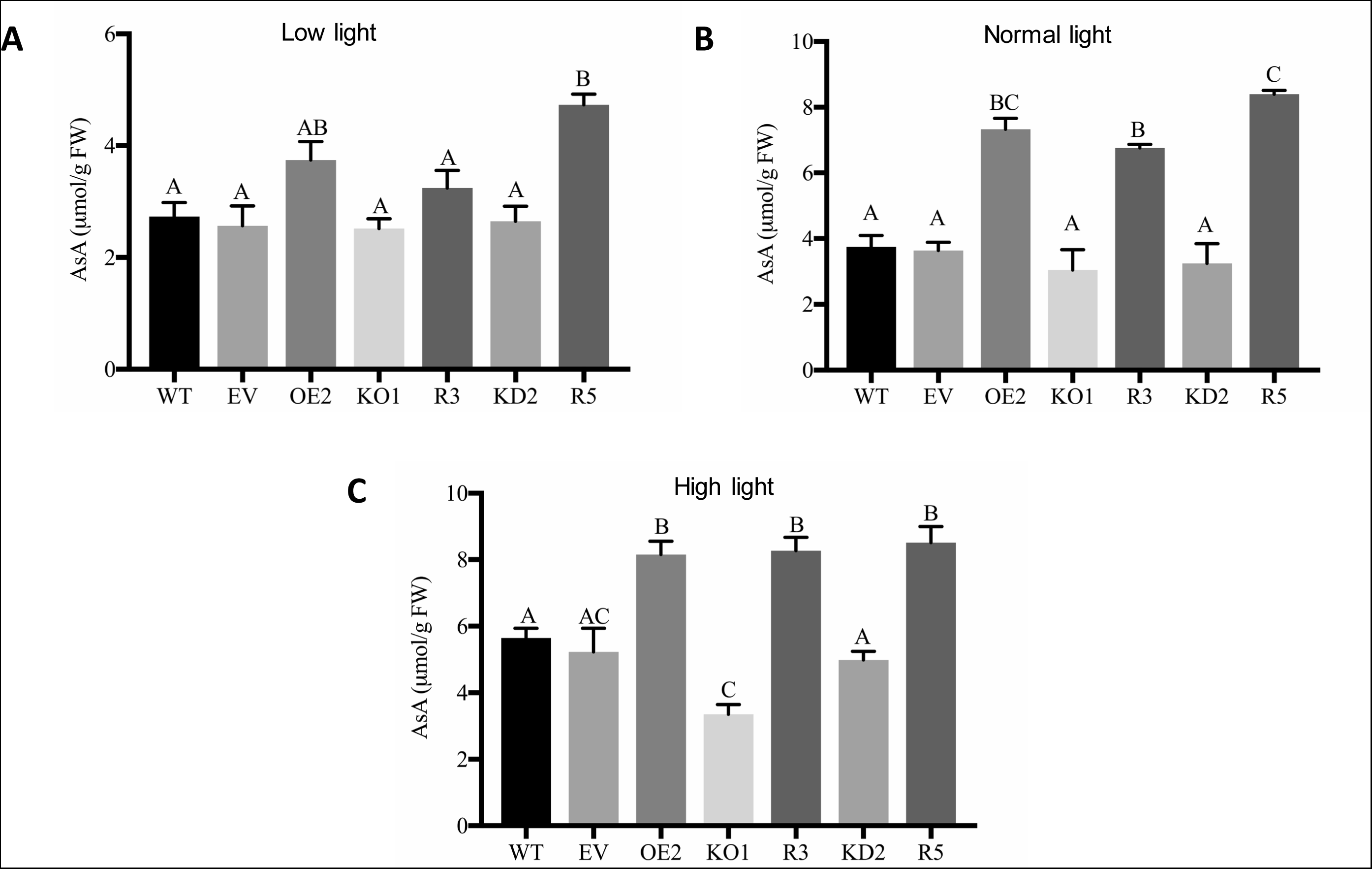
Total foliar As A levels of *At*GNL lines growing under (A) low, (B) normal, and (C) high light conditions. Each line was compared to the control (WT), analyzed by one-way ANOVA using Tukey’s posthoc test for multiple comparisons at 0.05 significance level. Significant differences are indicated by different letters. WT: wild type, EV: empty vector, OE: over-expresser, KO: knockout, R: restored. Values are means ± SEM (n=15).

*In planta* chlorophyll fluorescence measured with the fluorescence camera can serve as an indicator of whether the plants are under stress. High fluorescence in plants is opposite of high photosynthetic efficiency (Lichtenthaler, 1988). In Figure 11, the FLUO camera gave us a measure of relative chlorophyll fluorescence in *At*GNL lines. The knockout KO1 line showed high fluorescence compared to the other lines under high light. This knockout line has a lower As A level in the leaves, lower biomass, and project leaf area, and higher fluorescence compared with the rest of the lines. Overall these results show that the *At*GNL enzyme is essential to support normal AsA content in leaves and normal growth and development in Arabidopsis.

**Figure 11.**
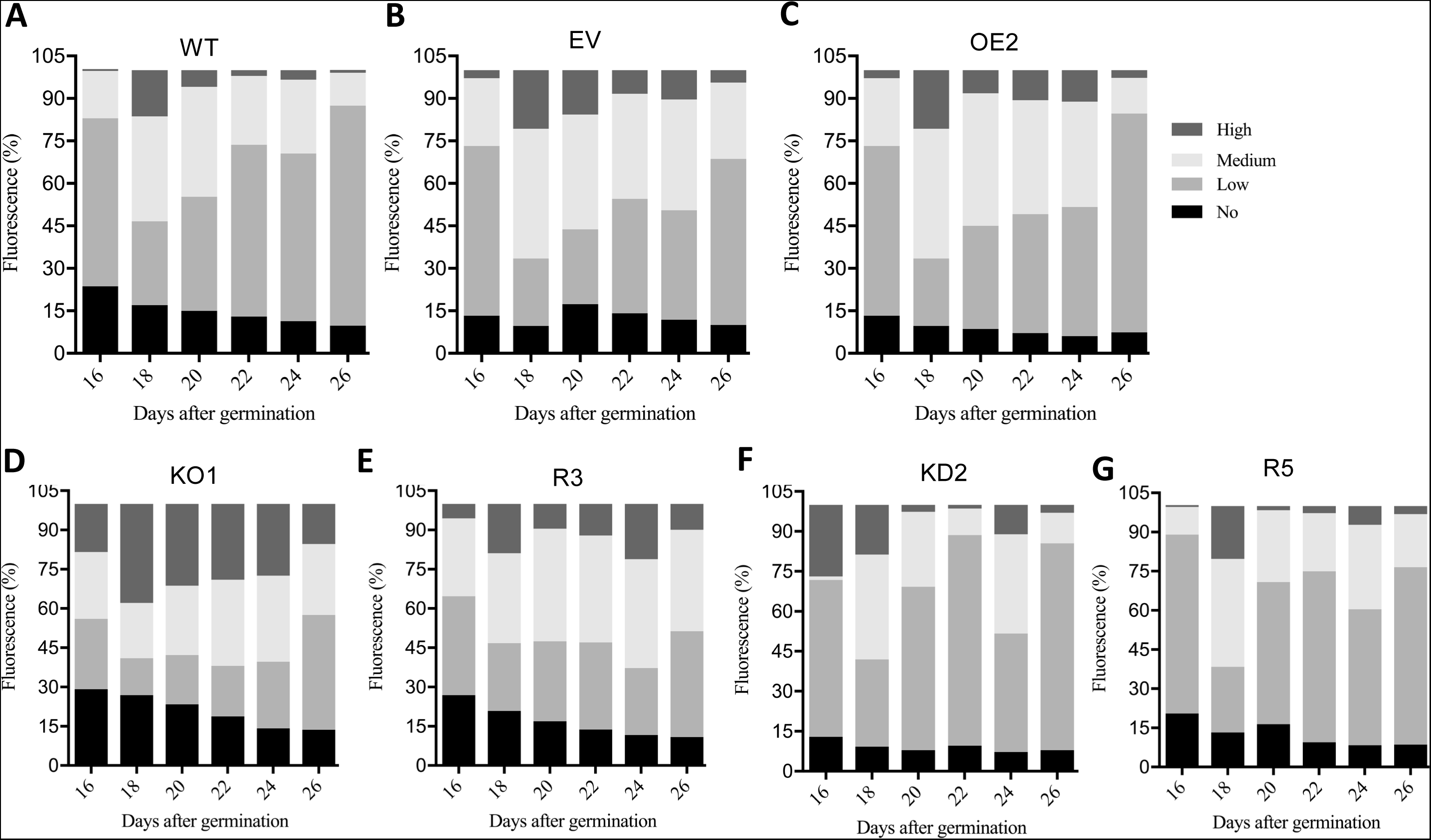
Chlorophyll fluorescence patterns of *At*GNL lines. Relative *in planta* chlorophyll content measured with the FLUO camera. WT: wild type, EV: empty vector, OE: over-expresser, KO: knockout, R: restored. Bars represent means (n=15).

### Photosynthetic Efficiency of *At*GNL Lines Under Low and Normal Light Conditions

Photosynthetic efficiency is the fraction of the light that plants obtain from the sun to convert into energy during photosynthesis. Photosynthetic efficiency was analyzed for *At*GNL lines growing under low and normal light conditions. Under normal light conditions there was no penalty in the photosynthetic efficiency of plants lacking *At*GNL expression (Figure 12). In contrast, the over-expressers and restored lines displayed enhanced efficiency indicating a positive impact on photosynthesis due to higher *At*GNL expression. When plants were grown under low light, results were quite different. In this case a lower photosynthetic efficiency was detected in EV and KO1 compared to the WT. This indicates a penalty in photosynthetic efficiency due to lack of *At*GNL expression. In addition to photosynthetic efficiency linear electron flow (LEF) was also measured. The linear electron flow rate has a direct correlation to photosynthetic efficiency. LEF facilitates the movement of H^+^ ions across the thylakoid membrane to create an electrochemical gradient that is used by ATP-synthase to produce energy (ATP). Figure 12 shows that under low light conditions both knockouts had lower LEF values than the controls, while KO1 was the line with the worst performance under normal light conditions. These data highlight the importance of the *At*GNL enzyme for efficiency ATP production in the chloroplasts. Plants exhibit phenotypic plasticity and respond to differences in environmental conditions by acclimation. In a recent study, Arabidopsis plants grown under field conditions were compared with plants grown indoors. Indoor-grown plants had larger leaves, modified leaf shapes and longer petioles and less NPQt, while field-grown plans had a high capacity to perform state transitions (Mishra et al., 2012). If photosynthesis is inefficient, excess light energy is dissipated as heat to avoid damaging the photosynthetic apparatus. When plants are under abiotic stress, such as low light intensity, the photosynthetic efficiency and the NPQt are opposite. Our results indicate that the KO1 line had high NPQt, indicating inefficient photosynthetic at both low and normal light conditions (Figure 12). Statistical analysis indicates the KO-1 line had a highly significant difference compared to the WT. Overall, our results showed that the over-expressers, restored lines, wild type, and empty vector lines had higher values of photosynthetic efficiency and LEF compared with KO lines under normal and low light conditions, while the NPQt values were opposite with the KO1 having the highest value. These result show that *At*GNL expression is essential to maintain high photosynthetic efficiency, high electron flow to make ATP (high LEF) and less loss of energy in the form of heat (NPQt).

**Figure 12.**
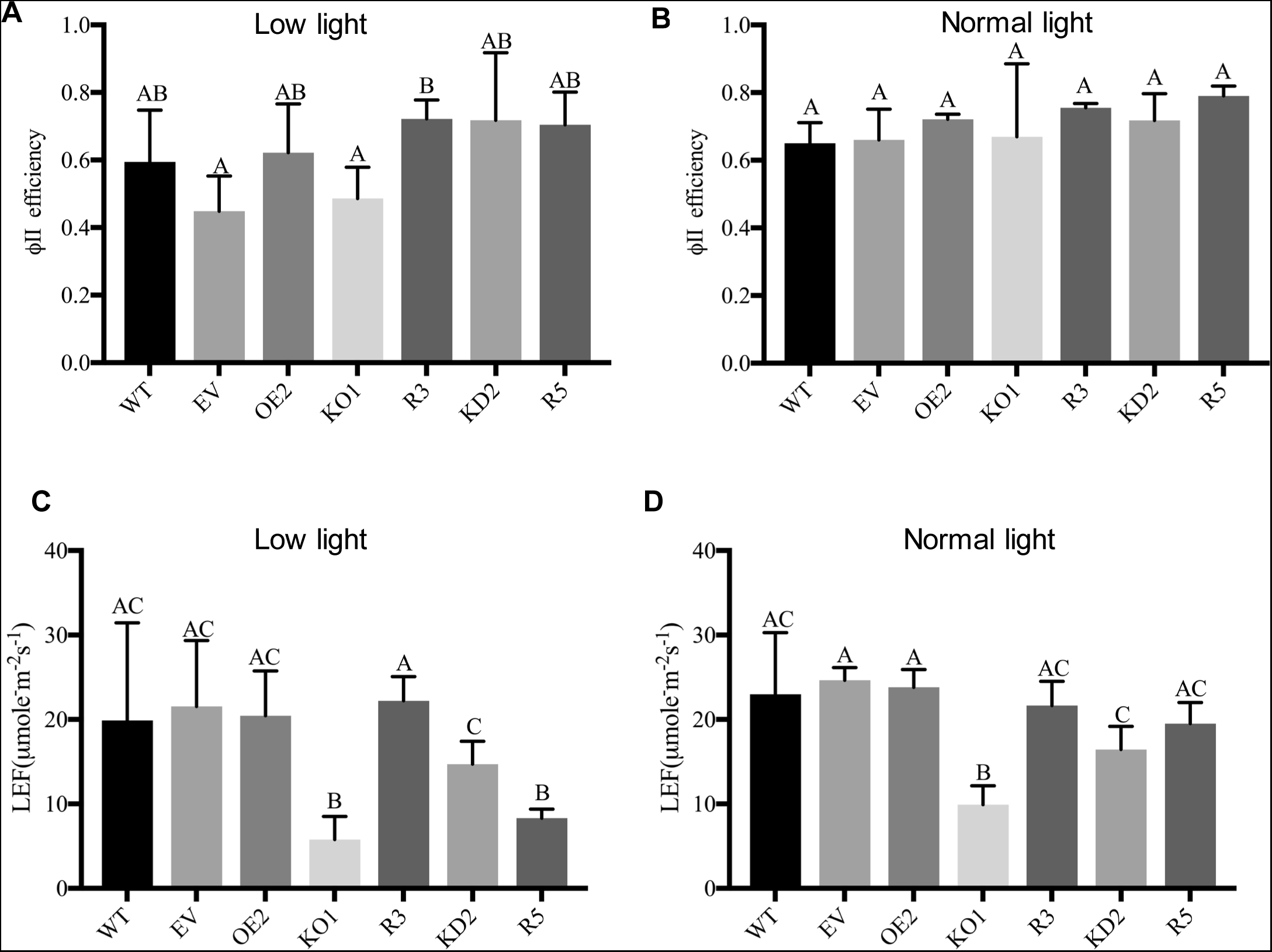
Photosystem II efficiency of *At*GNL lines grown under (A) low and (B) normal light conditions. Linear electro flow of *At*GNL lines grown under (C) low and (D) normal light conditions. Each line was compared to the control (WT), analyzed by one-way ANOVA using Tukey’s posthoc test for multiple comparisons at 0.05 significance level. Significant differences were indicated by different letters. WT: wild type, EV: empty vector, OE: over-expresser, KO: knockout, R: restored. Values are means ± SEM (n=10).

### Seed Yield of *At*GNL Lines Under Normal Light

In order to study the effect of light intensity on seed production, 15 plants of each line were grown in the greenhouse under medium light condition. Seeds were collected and counted. As illustrated in Figure 13, AtGNL constitutive expression had a dramatic effect on seed yield, as indicated by the 58.4% seed yield increase in OE2, and the 3.2-fold decrease in seed yield in KO1 compared to WT and EV controls.

**Figure 13.**
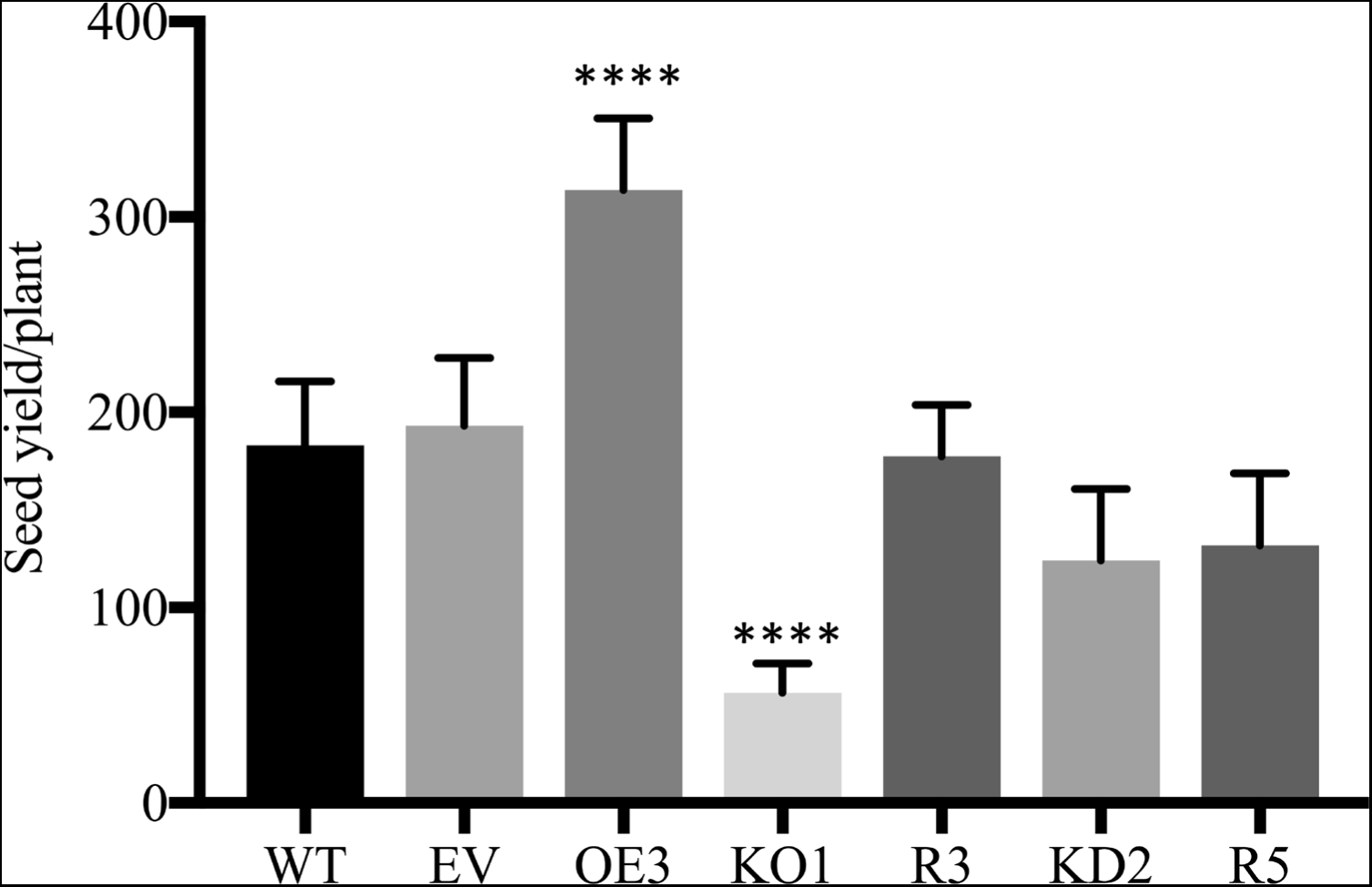
Seed yield of *At*GNL lines grown under normal light conditions. Values are means ± SEM (n=15). **** indicates significant differences at p˂0.0001 at 0.05 according to student’s t test compared means between WT control and other genotypes.

### Expression of *At*GNL Using the *GUS* Reporter Gene

In order to examine the expression of the *At*GNL within tissues, 10 transgenic plants expressing GUS driven by the *At*GNL promoter (p*At*1g56500: pCAMBIA1305.1) and empty vector pCAMBIA1305.1 (control) were generated. In the empty vector the *GUS-PLUS* gene is under the control of the 35S constitutive promoter. *At*GNL, empty vector, and WT plants were treated with the X-Gluc substrate. As shown in Figure 14, GUS activity was evident in plants expressing the *At*GNL promoter in all developmental stages from cotyledons to roots, although much less staining was observed in 4-day-old seedlings compared with controls. The oldest seedlings stained most intensely, especially at the leaf tips and margins. Our results show that the *At*GNL is expressed in the whole plant and at all developmental stages, indicating that the GNL enzyme is important in the physiological development of the plant from beginning to maturity.

**Figure 14.**
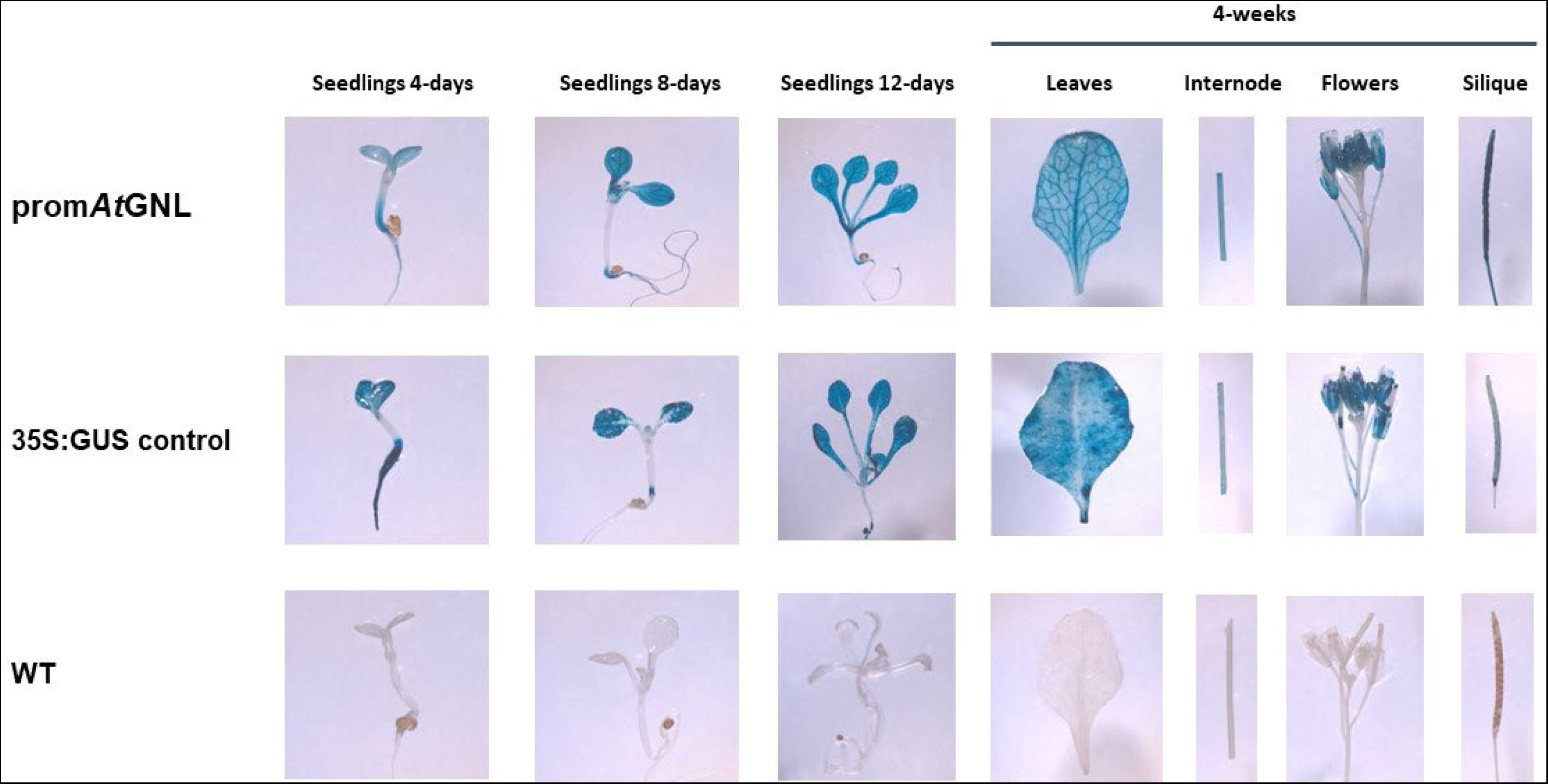
Temporal and spatial expression of *At*GNL using the *GUS-PLUS* reporter gene. Controls in the experiment were lines transformed with the empty vector where the *GUS-PLUS* gene is under the control of the 35S promoter (35S:GUS) and untransformed plants (wild type, WT).

### Phylogeny

The search for sequences homologous to *At*GNL (*At*1g56500) resulted in the deployment of 100 sequences from different dicotyledonous plants; only the sequences from *Arabidopsis*, *Brassica napus*, *Citrus sinensis*, *Gossypium hirsitum*, *Jatropha curcas*, were included in the phylogenetic tree (Figure 15). Those sequences showed a high similarity to the *At*GNL sequence (between 70 to 100% with BLASTP). To compare phylogenetically, those sequences were aligned with nine homologous sequences to regucalcin (senescence marker protein-30; RGN) from human, mouse, rat, and several bacteria. As illustrated GulLO and GNL gene families form distinct clades (groups) containing smaller subclades specific to dicots, Arabidopsis and animals. The sequences *Arabidopsis lyrata*, XP002891968.1 and *B. napus*, XP013696734 showed to highest level of sequence similarity to *At*1g56500.

**Figure 15.**
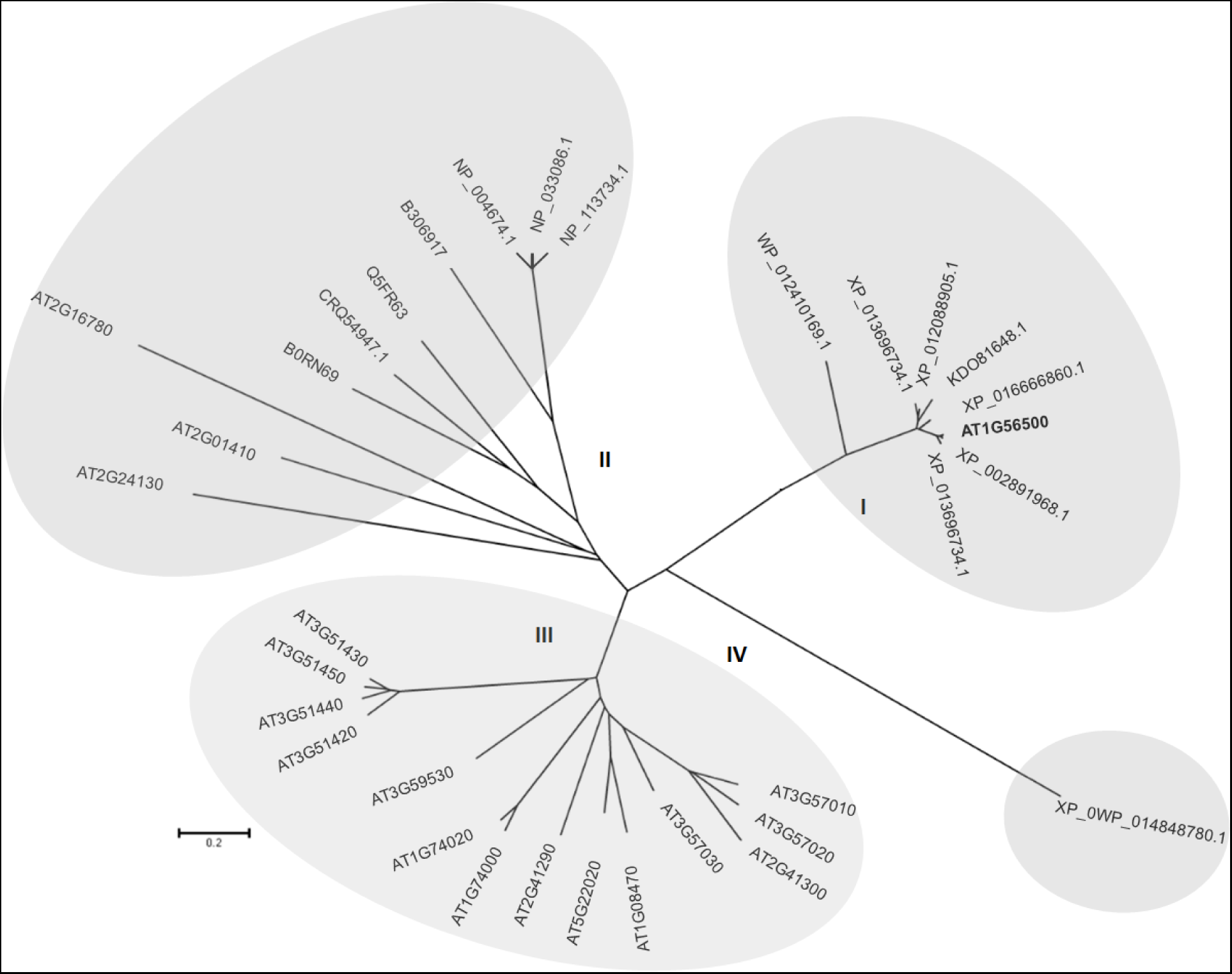
Phylogenetic tree for GNLs in plants and animals. The unrooted phylogenetic tree was constructed using MEGA version 5 using Neighbor-joining method and a bootstrap test with 1000 iterations. The tree was built with the multiple sequence alignment of the full-length, deduced protein sequences. The phylogenetic tree shows 18 putative GNL candidate genes in *A. thaliana* clustered in groups I and III (At1g08470, At1g56500, At1g74000, At1g74020, At2g01410, At2g16780, At2g24130, At2g41290, At2g41300, At3g51420, At3g51430, At3g51440, At3g51450, At3g57010, At3g57020, At3g57030, At3g59530, At5g22020) and six homologs sequences from different plan species clustered in group I (GenBank accession numbers are as follow: *B. napus*, XP013696734.1; *J. curcas*, XP012088905.1; *C. sinensis*, KDO816481; *G. hirsutum*, XP016666860.1; *A. lyrata*, XP002891968.1; as well characterized GNLs from rat (*R. norvegicus*, NP113734.1), human (*H. sapiens*, NP004674.1) and bacteria (*N. punctiform*, WP012410169.1; *Z. mobile*, WP014848780.1; *P. aeruginosa*, CRQ54947.1; *G. oxydans*, Q5FR63 and *X. campestris*, BORN69), groups I, II and IV.

## DISCUSSION

### Characterization of the *At*GNL Functional Enzyme

The protein encoded by At1g56500 was previously characterized by Brooks et al., 2013. They named it “*soq1*”. They described that this protein possesses a thioredoxin like domain found in thioredoxin-like domain (Trx-like domain), a lumen associated β-propeller domain, and a chloroplast stroma associated haloacid dehalogenase domain (HAD). In this work we propose that this protein has a moonlighting GNL activity and is involved in AsA biosynthesis. We demonstrated that the AtGNL enzyme is located in chloroplasts (Figure 4).

In our work the lactonase activity was assayed *in vitro* based on the decrease in absorbance (405 nm) of the *p*-nitrophenol pH indicator that resulted from the enzymatic opening of the lactone ring when D-glucono-σ-lactone (D-GuIL) was used as substrate in presence of the *At*GNL as previously described (Hucho and Wallenfels, 1972). Ogawa et al., (2002) reported that the GNL enzyme from *A. niger* had higher activity at 30°C, while the activity of the GNL from *P. aeruginosa* is optimal at 24°C (Tarighi et al., 2008), which are similar to the *At*GNL. Kondo et al., (2006), reported that the activity of the rat GNL was highest at pH 6.4 while Tarighi et al., (2008) demonstrated that the optimal activity of the *P. aeruginosa* GNL was at pH 7.2. In contrast, in this study the *A. thaliana* GNL enzyme had a higher activity at pH 6.0, and the activity decreased by 4-fold when pH was increased to 6.3 (Figure 5B). To assess if the *At*GNL activity preferred a particular divalent ion, various cofactors were tested. Ishikawa et al., (2008) reported that the GNL enzyme from *E. gracilis* had a higher activity using ZnCl_2_ as a cofactor and that this activity decreased around 4-fold when changed to MnCl_2_.

### Molecular and Phenotypic Characterization of *At*GNL Lines Under Light Stress

The *AtGNL* gene is significantly overexpressed in *At*GNL lines and restored lines (Figure 8). *At*GNL lines and restored lines have increased AsA compared to WT, KO1 and KD1 lines under normal light and light stress conditions (Figure 10A-C). The projected leaf area of over-expressers and restored lines was larger than their controls, with KO1 being the worst performer under low light, normal light and high light conditions (Figure 9). High ascorbate *A. thaliana* have elevated auxin and auxin-related cell elongation, increased intracellular glucose level, and enhanced photosynthesis, which are the reasons behind enhanced biomass in high ascorbate lines (Nepal et al., 2019). Low ascorbate lines (*vtc*1-1, *vtc*2-1, and *vtc*3) have lower AsA and are sensitive to heat and light stress (Pavet et al., 2005; Conklin et al., 2013). Additionally, low ascorbate lines (*vtc* mutants) are sensitive to oxidative and osmotic stresses (Cho et al., 2016). Under light stress and heat stress condition, low ascorbate *A. thaliana vtc*3 mutant have lower GalDH activity and AsA and are sensitive to light stress (Conklin et al., 2013). In contrast, high As A lines are tolerant to abiotic stresses. *A. thaliana* MIOX4 expressers are tolerant to heat stress, light stress, cold stress, and environmental pollutants (Tóth et al., 2011; Lisko et al., 2013). *Nicotiana tabacum* over-expressing monodehydroascorbate oxidase and dehydroascorbate oxidase are tolerant to methyl viologen stress, salt stress, cold stress, and oxidative stress indicated by increased growth and chlorophyll content (Eltelib et al., 2012; Kwon et al., 2003). High As A plants have enhanced biomass under salt stress (Cai et al., 2016; Eltelib et al., 2012; Hemavathi et al., 2009; Liu et al., 2015; Zhang et al., 2015), under cold stress (Cai et al., 2014), oxidative stress (Kwon et al., 2003), and light stress (Chen et al., 2015). Ascorbate is abundant in all actively growing and metabolically active plant tissues. A gradient of AsA concentration has been observed in different compartment of foliar tissue. In the absence of stress and low light conditions (150 μmol photons/m^2^/s), AsA concentration varies between cell compartments of leaf tissues as peroxisome (23 m M), cytosol (22 m M), nuclei (16 m M), chloroplasts (11 m M), mitochondria (10 m M), and vacuoles (2 m M) (De-Tullio and Arrigoni 2003; Zechmann 2018). When 2-week-old *A. thaliana* plants were grown under a 16 h photoperiod and 100 μmol photons/m^2^/s^1^ light intensity were transferred to dark conditions, the foliar As A content was significantly reduced, whereas when the plants were exposed to continuous light the AsA level in leaves increased by 171% (Yabuta et al., 2007). Similarly, shading of *Solanum lycopersicum* for 7 days reduced the As A and DHA in leaves by 50%, and a 10% reduction of As A and DHA was observed in fruits compared to the unshaded region (Massot et al., 2012). Low ascorbate lines (*vtc* 1-1, *vtc*2-1, and *vtc*3) have lower AsA and are sensitive to heat and light stresses (Pavet et al., 2005; Conklin et al., 2013). Additionally, these mutants are sensitive to oxidative and osmotic stresses (Cho et al., 2016). Under light stress and heat stress condition, *vtc*3 mutants have lower GalDH activity and As A and are sensitive to light stress (Conklin et al., 2013). Similarly, under heat and light stresses *A. thaliana vtc*2-3 mutants have lower photosynthetic efficiency and higher NPQt (Tóth et al., 2011).

### Photosynthetic Efficiency of *At*GNL Lines Under Low and Normal Light Conditions

Linear electro flow is lower in the KO1 line compared to other lines under normal and low light conditions (Figure 12C-D). Similarly, photosystem II efficiency is relatively lower in KO1 under low light condition (Figure 12A). Under heat and light stress *A. thaliana vtc*2-3 mutant have lower photosynthetic efficiency and higher NPQt and are sensitive to heat and light stress (Tóth et al., 2011). *Lycopersicum esculentum* overexpressing monodehydroascorbate reductase have increased net photosynthetic rate, and chlorophyll fluorescence under heat stress and cold stress (Li et al., 2010). High AsA *N. tabacum* have increased photosynthesis under ozone stress, drought stress, and salt stress (Eltayeb et al., 2006). High As A plants have increased chlorophyll content under oxidative stress, cold stress, and salt stress (Kwon et al., 2003). Similarly, *N. tabacum* overexpressing dehydroascorbate reductase have increased transpiration, and stomatal conductance under ozone stress (Chen and Gallie 2005). The photosynthetic rate is higher in monodehydroascorbate reductase overexpresser *L. esculentum* under heat stress and cold stress (Li et al., 2010). *A. thaliana* overexpressing dehydroascorbate reductase have enhanced stomatal aperture under to light, heat, and oxidative stress (Wang et al., 2010). Chlorophyll content is higher in high As A plants under drought stress (Eltayeb et al., 2011), salt stress (Cai et al., 2016; Li et al., 2012; Qin et al., 2015, Zhang et al., 2012), and light stress (Hu et al., 2016).

## CONCLUSIONS

The present study describes the characterization of an *Arabidopsis* gluconolactonase (*At*1g56500, *At*GNL), the third enzyme in the *MI* pathway to As A. This enzyme uses D-glucono-σ-lactone as a substrate *in vitro*. Analysis of *At*GNL *in planta* showed that *gnl* knockout (KO1) have low As A, grow slower, accumulate less biomass and are sensitive to light stress *in vivo*, indicating the importance of this enzyme to sustain effective plant growth and health. To our knowledge, this constitutes the first characterization of a plant GNL and is also the first As A biosynthetic enzyme known to function in the chloroplasts. The *At*GNL is active *in vitro* after purification via nickel column chromatography. The *At*GNL has a specific activity of 10.54 µmol/min/mg, slightly higher compared to the GNL from *G. oxydans*.

The function of the *At*GNL *in planta* was evident in the experiments where lines with normal (WT), lower (*gnl*: KO, KD), and higher expression (OE, R) were compared. The over-expressers and restored lines grew faster, accumulated more biomass, had higher seed yield and were more tolerant to light stress compared to their respective controls. Overall, we conclude that *At*GNL is important for plant development and protection of the plant from light stress. Photosynthetic efficiency is an important parameter that is directly correlated with proper plant development and growth. Results shown here demonstrate that the over-expresser and restored lines had higher photosynthetic efficiency than the controls growing under normal and low light conditions. The same pattern was observed linear electron flow (LEF). In addition, when non-photochemical quenching was analyzed the *gnl* knockout (KO1) performed significantly poorer compared to the other lines, indicating the importance of this enzyme to sustain proper function of the photosynthetic machinery.

Spatial and temporal expression analysis of the *At*GNL promoter fused to the GUS reporter showed *At*GNL is found throughout the plant, from early developmental to the development of siliques, indicating that the *At*GNL is expressed constitutively. Taken together these results indicate that AtGNL is an active enzyme involved in ascorbate metabolism and in maintaining proper function of the photosynthetic machinery in plants experiencing light stress. Photosynthetic efficiency has been shown to have a high correlation with global crop productivity (Murchie et al., 2015). It would be interesting to study the effect of over-expressing this *At*GNLs in crops of agricultural importance.

## MATERIALS AND METHODS

### Plant Material and Growth Conditions

*Arabidopsis thaliana* ecotype Columbia wild type (Col-0, Stock # CS-60000), SALK_026172, and SALK_011623 were obtained from the Arabidopsis Biological Resource Center (ABRC, Columbus, OH). Seeds were sterilized with 70% (v/v) ethanol followed by 50% (v/v) sodium hypochlorite containing 0.05% (v/v) Tween-20. Next, seeds were washed with sterile water. Seeds were transferred to petri dishes containing MS medium: MS salts, sucrose 3% (w/v) at pH 5.6 and supplemented with 0.04% (w/v) MgSO_4_.7H_2_O. After stratification at 4°C for 3 days, plates were transferred to a growth chamber and incubated at 23°C day, 65% humidity, 16:8 h photoperiod and 200 µmol/m^2^/s light intensity. Finally, robust seedlings were transferred to PM-15-13 AIS MIX Arabidopsis soil (Lehle-Seeds, Round, TX) and grown to maturity under the above cited conditions.

*Nicotiana benthamiana* seeds were obtained from Dr. Sue Tolin at the Department of Plant Pathology, Physiology and Weed Science at Virginia Polytechnic Institute and State University (Blacksburg, VA). Seeds were sown in 4.5-inch pots containing Pro-mix BX soil (Premier Horticulture) with Osmocote 14-14-14 fertilizer (Scotts). Vermiculite was overlaid on top of the seeds. Plants were grown in an environmental control chamber with the following conditions: 25°C (day)/21°C (night) temperature, 65% relative humidity, 16:8 h photoperiod, and 150 µmol/m^2^/s light intensity.

### Gene Constructs and Plant Transformation

Two gene constructs were used in this work. A first one where the *At*GNL ORF is under the control of the 35S promoter and the tobacco etch virus (TEV) enhancer (*At156500-6XHIS:pBIB-Kan*), and a second construct where the 1,000 bp promoter region of the *At*GNL fragment preceding of the ATG, was cloned and fused to the *GUS* reporter gene (*pAtGNL:pCAMBIA1305.1*) (Figure 2).

Stable Arabidopsis transformation was achieved by the floral dip method (Clough and Bent, 1998) using *Agrobacterium tumefaciens* GV3101 strain carrying the construct of interest. The transgenic status of the Arabidopsis lines was established via PCR using genomic cDNA as a template.

### Transient Protein Expression

Transient AtGNL expression was performed in *Nicotiana benthamiana*. Five week old tobacco plants were vacuum infiltrated with the *AtGNL-6XHIS:pBIB-Kan* construct as described by Medrano et al., 2009. Leaves were harvested 48 h post infiltration, frozen immediately in liquid nitrogen and stored at −80°C until further processing. The expression of *At*GNL was confirmed via Western blot as follows. Crude extracts were made by grinding frozen tissue in presence of two volumes of SDS buffer containing: 150 m M Tris-HCl pH 6.8, 5 m M EDTA pH 8.0, 30% (v/v) glycerol, 6% (w/v) SDS. Proteins were separated via SDS-PAGE on 10% precast mini-gels (Expedeon) with a Tris-MOPS buffer. Recombinant *At*GNL-6XHIS was detected using an anti-HIS (C-term)/AP antibody at a 1:2,000 v/v dilution (Invitrogen) and CDP-start, a chemiluminescent substrate for alkaline phosphatase detection (Roche Diagnostics).

### Protein Purification

Recombinant *At*GNL protein was purified from *N. benthamiana* leaves. Five grams of leaves were pulverized in liquid nitrogen and protein was extracted with 10 mL of buffer A containing: 75 m M sodium phosphate dibasic, 25 m M sodium phosphate monobasic, 150 m M NaCl, 10 m M sodium metabisulfite, and 0.6% (v/v) protease inhibitor cocktail, pH 7.4. The extract was then centrifuged at 13,000 x g for 15 min. The resulting supernatant was loaded onto a nickel affinity column (HIS60 Ni Superflow) and incubated for 1 h at 4°C. Then, the column was washed with 50 m M sodium phosphate pH 7.4, 300 m M NaCl, 40 m M imidazole buffer and the bound proteins were eluted with 250 m M of imidazole. The eluate was concentrated using an AMICON^®^ 30K ultra centrifugal filter (Millipore). Total soluble protein concentration was estimated by the Bradford method (Bradford, 1976) using Coomassie blue G-250 dye (Thermo Scientific) and bovine serum albumin (Pierce) as a standard. Protein fractions from the purification procedure were separated by SDS-PAGE and the *At*GNL was detected by Western blot and silver staining using the Pierce® Silver Stain Kit (Thermo Scientific) following the manufacturer’s instructions.

### Enzyme Assays

The optimum concentration for enzyme activity was determined to be 30 µg per reaction. The lactonase activity was assayed *in vitro* based on the decrease in absorbance (405 nm) of the *p*-nitrophenol pH indicator as previously described (Ishikawa et al., 2008). Briefly, one mL of the reaction contained: 10 m M PIPES pH 6.5, 5 m M D-glucono-δ-lactone, 75 µM MnCl_2_, 2.5 m M *p*-nitrophenol, and an aliquot of the purified enzyme. An equal amount of boiled enzyme was used as control for these experiments. Multiple substrates were tested in the lactonase assay: D-glucono-δ-lactone (D-GuIL), L-galactono-ɤ-lactone (L-GaIL), L-galactonic acid (L-GaIA), L-gulono-ɤ-lactone (L-GuIL), and L-gulonic acid (L-GuIA). The L-GaIA and L-GuIA were prepared by the hydrolysis of the L-GaIL and L-GuIL, respectively. For hydrolysis 20 mL of 0.3 M NaOH were added to 100 µL of the 10 m M L-GaIL or L-GuIL, the mixture was vigorously agitated by vortexing for 20 s, and 20 µL of the 0.3 M HCl were added to neutralize the solution (Ishikawa et al., 2008). The enzyme kinetic experiments reaction was monitored at different substrate concentrations. Enzyme kinetic analysis was done using GraphPad Prism 6.2 (GraphPad Software).

### RT-qPCR Analysis

Real time quantitative polymerase chain reaction (RT-qPCR) were performed for validation of *AtGNL* expression using reference genes previously published and following MIQE guidelines (Czechowski et al., 2005; Bustin et al., 2009). Three biological replicates of the leaf samples were collected when plants that completed the vegetative growth stage (stage 5.0 according to Boyes et al., 2001) in normal light treatment. Total RNA was purified using Purelink™ RNA mini kit (Ambion) following the manufacturer’s instructions. Residual DNA in the sample was removed using DNA-*free*™ DNA removal kit (Invitrogen). The RNA quality and quantity was assessed using the Experion™ RNA HighSens chips (BioRad). The cDNA was prepared by following manufacture’s instruction (iScript select cDNA synthesis kit, BioRad). The GNL primers (GTTTGGAGA CAATGATGGCGT/TCCCGTTCCAGCGAGAGTAAC) were validated as suggested by Bustin et al., 2009. Real time quantitative PCR was performed following manufacturer’s instruction (SsoFast™ EvaGreen^®^ Supermix, BioRad) in a CFX-384™ real-time system (BioRad). Normalization of transcripts counts was done using the *ACTIN2* (GTATCGCTGACCGTATGAG/ CTGCTGGAATGTGCT GAG) and *EF1-α* (GGTGGTGGCATC CATCTTGTTACA/GGTGGTGGCATC CATCTTGTTACA) reference genes. The relative expression was calculated using ΔΔCq (Pfaffl 2004). Three biological replicates and two technical replicates for each genotype were used for RT-qPCR. Student’s independent *t*-test was performed at 0.05 significance level.

### Ascorbate Measurements

*In planta* AsA concentration in Arabidopsis changes throughout the day as well as during development (Tamaoki et al., 2003; Zhang et al., 2009). Fifty mg of leaf tissue were collected between 9:00 and 11:00 am. Tissue was frozen immediately in liquid nitrogen and stored at −80°C until analyzed. Reduced, oxidized, and total As A were measured using 96-well plate format as described by Haroldsen et al., (2011). Frozen tissue was pulverized in 6% (w/v) *meta*-phosphoric acid and centrifuged at 13,000 rpm for 15 min. Reduced As A was determined by measuring the decrease on absorbance at 265 nm after addition of 0.5 unit of ascorbate oxidase to 300 µL of the reaction medium containing the plant extract and 100 m M phosphate buffer at pH 6.9. Oxidized ascorbate was measured in 300 µL reaction mixture with 10 µL of 40 m M dithiothreitol (DTT) after incubation in the dark for 20 min at room temperature. Calculations were made based on a standard curve made with pure L-ascorbic acid run in parallel.

### Light Stress Assays

For the analysis of light stress tolerance, transgenic homozygous Arabidopsis seeds were surface-sterilized with bleach and place in petri dishes with MS medium for 10 days. Seedlings were transferred to PM-15-13 AIS MIX Arabidopsis soil (Lehle-Seeds) in Quick15 pots. Trays were covered with a dome for one week in grow chamber, then plants were transferred to the greenhouse. Growth conditions were as follows: 22°C - 26°C temperature, 16:8 h photoperiod, 55% humidity and three different light conditions: low (35-110 µmol/m^2^/s), normal (110-350 µmol/m^2^/s), and high light (350-700 µmol/m^2^/s). Light intensity was recorded four times per day (9:00 am, 12:00 pm, 3:00 pm, and 6:00 pm) to cover the entire sunlight period. This work was performed during March 12 - 30, 2015 in Jonesboro, AR, USA (latitude 29.4889 and longitude −98.3987).

### High Throughput Phenotyping

The phenotype of Arabidopsis lines of interest grown under low, normal, and high light conditions was studied using a high throughput phenotyping platform (Scanalyzer HTS, Lemnatec, Germany) and the LemnaControl software (LemnaTec). This instrument is equipped with a robotic arm that holds visible (VIS, a.k.a RGB), fluorescence (FLUO), and near infrared (NIR) high-resolution cameras. The cameras in the system are as follows: VIS camera, piA2400-17gc CCD (Basier) with resolution of 2454 x 2056 pixels; FLUO camera, scA1600-14gc CCD (Basler) with resolution 1624 x 1234 pixels; and NIR camera, Goldeye GIGE P-008 SWIR (Allied Vision Technologies) with resolution 320 x 256 pixels and with spectral sensitivity between 900 and 1700 nm. Images of *At*GNL lines were captured every two days between 16 days and 26 days after germination, to cover the full vegetative growth. Images were analyzed using the LemnaGrid Software. The analysis of the RGB images was done as previously described by Acosta-Gamboa et al., (2016) Multiple phenotypic parameters were calculated for each plant including: projected leaf area (cm^2^), convex hull area (cm^2^), caliper length (rosette diameter) and compactness (measure of the bushiness of the plant). From the RGB images the relative area of the plants displaying normal green color versus the area with detectable yellow color (chlorosis) was calculated. The analysis of the NIR images, was similar to the color classification of VIS camera, using the acquired gray-scale images, where high water content corresponds to darker tones while low water content corresponds to lighter gray tones. The software used this information to calculate the relative area with low, medium, and high-water content. The fluorescence camera acquired red-scale images and in this case the red tones were divided into four equidistant bins, and the software calculated the relative area with zero, low, medium, and high fluorescence. Quantitative data obtained from the images was exported as CSV files and analyzed in Excel.

### Photosynthetic Efficiency Measurements

Photosynthetic efficiency of photosystem II (Φ/II) and linear electron flow (LEF) of the Arabidopsis lines of interest growing under low, and normal light conditions were analyzed using a MultispeQ (Kulhgert et al., 2016). This is a hand-held device developed by the Kramer Laboratory at Michigan State University. Data was visualized in an Android tablet (Samsung Galaxy Tab 4) and analyzed in the PhotosynQ website (www.photosynq.org).

### Seed Yield Assessment

In order to study the effect of light intensity on seed production, fifteen plants of each *At*GNL lines were grown in the greenhouse under normal light condition (110-350 µmol/m^2^/s). Plants were harvest individually after the siliques (fruits) have completely browned and stored in paper bags. Dry plants were placed directly onto sieve and crushed using hands to remove all the seeds from the siliques. After sieving, the seeds were counted using the Elmor C3 Seed Counter (Elmor, Switzerland).

### Promoter *At*GNL:GUS Expression in *Arabidopsis thaliana*

To study the expression of the *At*1g56500 in different plant tissues, *A. thaliana* was transformed by the floral dip method (Clough and Bent, 1998) with *A. tumefaciens* GV3101 carrying the *pAtGNL:pCAMBIA1305.1* construct. A different set of plants was also transformed with bacteria carrying the empty vector control (pCAMBIA1305.1). The presence of the transgene of interest was established via PCR using gene specific primers, and genomic cDNA as a template. Explants (seedlings, leaves, flowers, and fruits) were cut from plants 4, 8, 12, and 30 days after germination. Next, explants were incubated in fresh and cold phosphate buffer pH 7.0 with 4% formaldehyde at room temperature for 30 min. The explants were washed several times with cold phosphate buffer for 1 h, the vacuum infiltrated with X-Gluc substrate solution containing: 1 mg 5-bromo-4-chloro-3-indolyl *β*-D-glucuronide in 100 µL of methanol, 1 mL 2X phosphate buffer, 20 µL 0.1 M potassium ferrocyanide, 20 µL 0.1 M potassium ferricyanide, 10 µL 10% (w/v) solution of Triton X-100, and 850 µL of water. Tissues were incubated in darkness at room temperature overnight until a distinct blue staining appeared. Finally, explants were incubated in 70% ethanol until the chlorophyll was removed. Photographs were taken with AxioCam MRc camera connected to a Stemi 2000-C stereo microscope (Zeiss).

### Phylogeny Analysis

The amino acid sequence for *At*1G56500 from the Arabidopsis Information Resource (TAIR: http://www.arabidopsis.org/) database sequence was used to search for homologs at NCBI (http://www.ncbi.nlm.nih.gov/) with BLASTP (Altschul et al., 1997). The homologous protein sequences were aligned using Multiple Sequence Comparison by Long-Expectation (MUSCLE) with the default parameters in Molecular Evolutionary Genetics Analysis MEGA5 (Kumar et al., 2008). A phylogenetic tree was constructed with MEGA5 using the neighbor joining method with 1,000 bootstrap replicates.

### Statistical Analysis

Data were analyzed using GraphPad prism v7.0 (https://www.graphpad.com). Independent student’s t-test and one way-ANOVA using Tukey’s post-hoc test for multiple comparisons were used to analyze the data at 0.05 significance level.

## Contributions

JPY did most of the experiments and wrote the initial draft of the manuscript

NN did RT-qPCR experiments, statistical analysis, assisted in figure and manuscript preparation

TKT demonstrated that AtGNL expressed in tobacco is localized in the chloroplast

SIA sequenced all DNA constructs and provided key expertise and training in protein purification

GAW did initial screening of Arabidopsis T-DNA lines and transformed Arabidopsis plants

AW did ascorbate measurements and light stress assays KMJ developed the phylogenetic tree

AL conceived the project, secured funding, identified putative GNLs in the Arabidopsis genome, design and supervised the experiments, and prepared the final version of the manuscript

## ACKNOWLEDGMENTS

This work was supported by a sub-award from the IDeA Networks of Biomedical Research Excellence INBRE program [National Center for Research Resources (5P20RR016460-11) and the National Institute of General Medical Sciences (8P20GM103429-11) from the National Institutes of Health] and by the Plant Imaging Consortium (NSF EPSCoR Track-2 Research Infrastructure Improvement Program Award IIA-1430427. We also thank funds provided by the Arkansas Biosciences Institute, the major research component of the Arkansas Tobacco Settlement Proceeds Act. AL thanks summer salary support from the AR INBRE program. JPYC and NN thank the Molecular Biosciences Graduate Program at Arkansas State University for stipend support. We thank Guillermo Trujillo-Lujan for making the AtGNL:pBIBKan and the pAtGNL:GUS constructs and Kim Lee for assistance with plant care.

## FIGURE LEGENDS

**Supplementary Figure 1.** Effects of co-factors and substrate on *At*GNL enzyme activity. (A) cofactors effect on *At*GNL activity. (B) effects of substrate (D-Glucono-σ-lactone) concentration on *At*GNL activity. Values are means ± SD.

**Supplementary Figure 2.** Experimental set up for studying the effect of light on the phenotype of *At*GNL lines. Light intensity was measures four times a day to cover the entire sunlight period.

