## SupplementalFigures for "Arabidopsis Gluconolactonase, the First Enzyme Involved in Ascorbate Biosynthesis Localized in the Chloroplast Protects Plants from Light Stress"

### Supplementary figures

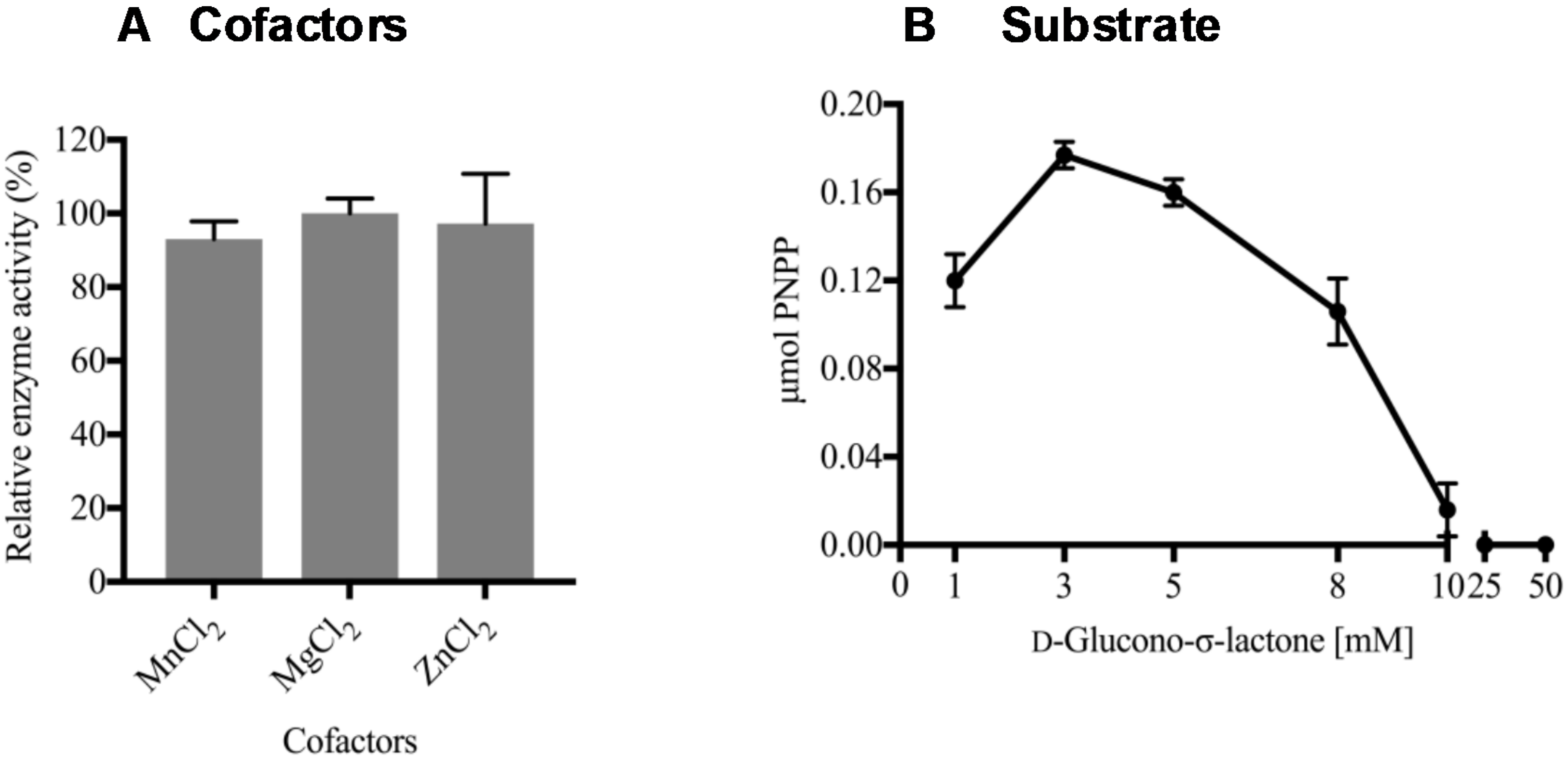


Supplementary Figure 1. Effects of co-factors and substrate on *At*GNL enzyme activity. (A) cofactors effect on *At*GNL activity. (B) effects of substrate (D-Glucono-σ-lactone) concentration on *At*GNL activity. Values are means ± SD.

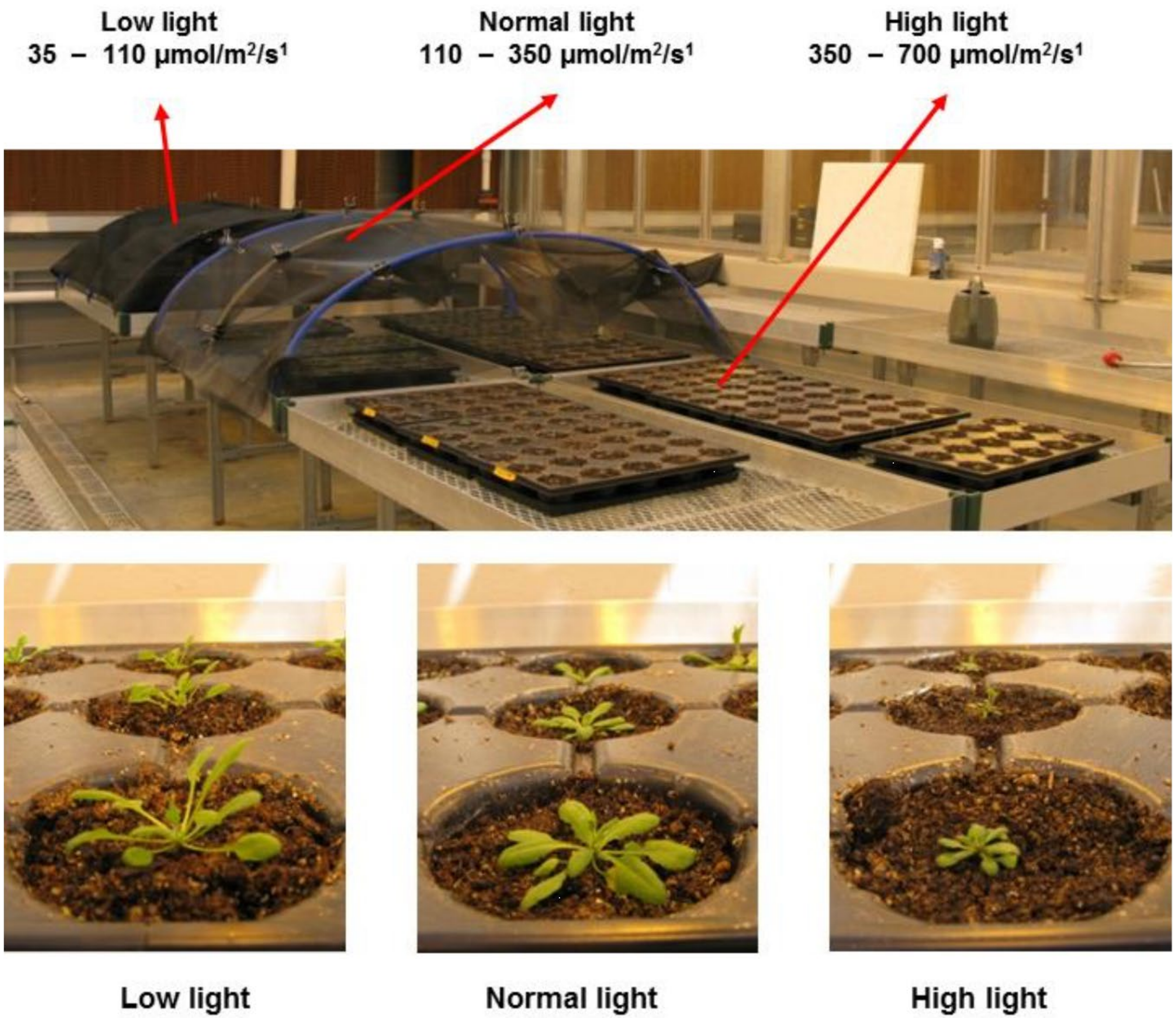


Supplementary Figure 2. Experimental set up for studying the effect of light on the phenotype of *At*GNL lines. Light intensity was measures four times a day to cover the entire sunlight period.
